# The SAS Chromatin-Remodeling Complex Mediates Inflorescence-Specific Chromatin Accessibility for Transcription Factor Binding

**DOI:** 10.1101/2024.02.20.581124

**Authors:** Jing Guo, Zhen-Zhen Liu, Yin-Na Su, Xin-Jian He

## Abstract

While the role of transcription factors in flower development is well understood, the impact of chromatin remodeling on this process remains largely unclear. We conducted a comprehensive analysis to investigate the coordination of the SAS, BAS, and MAS-type SWI/SNF chromatin-remodeling complexes with transcription factors to regulate chromatin accessibility and gene transcription during flower development in *Arabidopsis thaliana*. Our findings indicate that the SAS complex binds to numerous genes related to flower development and is responsible for establishing chromatin accessibility of these genes during flower development. In contrast, the BAS and MAS complexes exhibit minimal involvement in regulating the accessibility of these genes. The SAS-bound genomic regions and the SAS-dependent accessible regions are enriched with sites occupied by multiple MADS family transcription factors involved in flower development. Furthermore, we found that the SAS-dependent accessibility is indispensable for the genomic binding of the MADS transcription factor AP1 at these regions. This study highlights the dynamic role of the SAS complex in modulating the genomic binding of transcription factors during plant development.

**One-sentence summary:** The SAS-type SWI/SNF complex regulates Inflorescence-specific chromatin accessibility at distal promoter and upstream intergenic regions, thereby facilitating the binding of transcription factors involved in flower development.

## Introduction

Accurate gene expression in specific cells and developmental stages is crucial for the development of organisms. ATP-dependent chromatin-remodeling complexes play a vital role in facilitating the access of transcription factors and transcription machinery to DNA by utilizing the energy derived from ATP hydrolysis, thereby driving dynamic changes in gene expression. The SWI/SNF (Switch defective/Sucrose non fermentable) complex, originally discovered in yeast, was the first identified chromatin-remodeling complex (Cairns et al., 1994; Peterson et al., 1994). In *Drosophila melanogaster*, SWI/SNF components were found to be Trithorax group factors that counteract the repressive effect of Polycomb group proteins on the transcription of specific homeotic genes, ensuring the accurate expression of these homeotic genes at precise positions (Kennison and Tamkun, 1988; Tamkun et al., 1992). Moreover, extensive studies in mammals have underscored the significance of SWI/SNF complexes in maintaining pluripotency and driving cell fate determination during development (Ho and Crabtree, 2010; Bieluszewski et al., 2023). Mutations in SWI/SNF subunits in humans have been closely linked to various types of tumors and developmental disorders (Cenik and Shilatifard, 2020).

In *Arabidopsis thaliana*, the SWI/SNF-type chromatin-remodeling ATPases consist of BRAHMA (BRM), SPLAYED (SYD), and MINUSCULE1/2 (MINU1/2). Some of the Arabidopsis SWI/SNF components were found to overcome the Polycomb repression (Wu et al., 2012; Li et al., 2015; Li et al., 2016; Shu et al., 2021), suggesting an evolutionarily conserved function of SWI/SNF complexes in plants and metazoans. Recent studies utilizing affinity purification followed by mass spectrometry (AP-MS) analysis have identified three classes of SWI/SNF complexes in Arabidopsis: the BRM-associated SWI/SNF (BAS) complex, the SYD-associated SWI/SNF (SAS) complex, and the MINU1/2-associated SWI/SNF (MAS) complex. Each of these complexes is composed of 8-15 subunits, with some subunits being specific to a particular complex, while others are shared by two or three complexes (Guo et al., 2022; Fu et al., 2023). However, the precise roles of different Arabidopsis SWI/SNF complexes in specific developmental processes remain elusive.

The effects of SWI/SNF components on plant growth and development have been extensively studied. Single mutants of *brm* and *syd*, as well as the weak *minu1/2* double mutant, display pleiotropic developmental defects, indicating the crucial role of SWI/SNF complexes in plant development. Moreover, the combination of *brm* and *syd* mutations, along with the *minu1/2* double knockout mutations, leads to lethality, further emphasizing the importance of these complexes (Wagner and Meyerowitz, 2002; Farrona et al., 2004; Bezhani et al., 2007; Sang et al., 2012). Notably, the mutants of BAS, SAS, and MAS components exhibit both overlapping and distinct phenotypes throughout Arabidopsis development (Bezhani et al., 2007; Guo et al., 2022). In particular, SAS-specific mutants, such as *syd*, *swi3d*, and *sys1/2/3*, consistently show severe inflorescence developmental phenotypes, including defects in shoot meristem maintenance, splayed-opened sepals, dramatic homeotic transformation of floral organs, and complete sterility (Wagner and Meyerowitz, 2002; Kwon et al., 2005; Sarnowski et al., 2005; Guo et al., 2022). The BAS mutant *brm* also exhibits flower defects, including abnormal size and number of floral organs, homeotic transformation of the second and third flower whorls, fused filaments, and reduced fertility (Farrona et al., 2004; Hurtado et al., 2006). However, the floral homeotic defects in *brm* are generally milder compared to SAS-specific mutants (Guo et al., 2022). Additionally, other BAS-specific mutants, such as *swi3c*, *brip1/2*, and *brd1/2/13*, show weaker or nearly normal flower and silique phenotypes in comparison to *brm* (Archacki et al., 2009; Yu et al., 2020; Jaronczyk et al., 2021; Yu et al., 2021; Guo et al., 2022). Although MAS-specific mutants, including *minu1/2* and *pms2a/b*, have been reported to display abnormalities in the internal structures of flowers, shorter siliques, and reduced fertility, their sepals and petals develop normally (Sang et al., 2012; Diego-Martin et al., 2022; Guo et al., 2022). These studies suggest that SAS plays a more important role in flower development compared to BAS and MAS.

Flower development encompasses multiple stages, including floral induction, floral meristem formation, and floral organ development, which is a strictly controlled program regulated by a network of transcription factors, with the LEAFY (LFY) transcription factor and various MADS-box transcription factors playing crucial roles (Weigel, 1995). The genetic association between SWI/SNF components and flower development-related transcription factors was established over two decades ago when the *syd* mutation was found to enhance the floral homeotic defects in the weak *lfy* mutant (Wagner and Meyerowitz, 2002; Bieluszewski et al., 2023). In *brm* and *syd*, the expression of several floral homeotic genes is downregulated (Hurtado et al., 2006; Wu et al., 2012; Lin et al., 2023). It has been reported that LFY and the MADS-box transcription factor SEP3 recruit BRM and SYD to counteract Polycomb repression at the floral homeotic genes *AP3* and *AGAMOUS* (*AG*) during flower development (Wu et al., 2012). The AUXIN RESPONSE FACTOR (ARF) MONOPTEROS (MP) was also reported to recruit BRM and SYD to increase the accessibility of its targets involved in flower formation upon auxin sensing (Wu et al., 2015). Additionally, the AP-MS results have demonstrated that BRM, SYD, and certain other SWI/SNF subunits are co-purified with multiple MADS transcription factors in flowers (Smaczniak et al., 2012), indicating interactions between SWI/SNF components and MADS transcription factors involved in flower development. A recent report has shown that LEAF AND FLOWER RELATED (LFR), a shared subunit of SAS and MAS, collaborates with the SAS catalytic subunit SYD to regulate the chromatin state and transcription of *AG* (Lin et al., 2023). These studies have mainly focused on the regulation of SWI/SNF components on specific genes associated with flower development. However, how the SWI/SNF complexes are coordinated with transcription factors to regulate gene transcription during flower development at the whole-genome level remains largely unclear.

In this study, we conducted a genome-wide analysis to compare the impact of BAS, SAS, and MAS complexes on chromatin accessibility and gene expression between seedlings and inflorescences. Our findings revealed that the SAS complex is responsible for establishing chromatin accessibility for a substantial subset of flower development-related genes in inflorescences as compared to seedlings. The SAS-bound genomic regions as well as the SAS-dependent accessible regions are enriched with genomic regions bound by MADS family transcription factors. The chromatin accessibility generated by SAS is required for the genomic binding of APETALA1 (AP1) at these regions. These results reveal the specific role of SAS in regulating the genomic binding of MADS transcription factors during flower development, providing insights into the molecular mechanism underlying the dynamic role of chromatin-remodeling complexes.

## Results

### SAS exhibits enhanced impact on gene expression in inflorescences relative to seedlings

We previously have determined the genome-wide effect of three classes of SWI/SNF complexes on chromatin accessibility and gene expression in the seedlings (Guo et al., 2022). However, whether and how the SWI/SNF complexes dynamically regulate chromatin accessibility and gene expression in different development processes are largely unclear. SAS mutants show several specific developmental phenotypes relative to BAS and MAS mutants (Guo et al., 2022), a prominent phenotype among which is the severe floral developmental defects (Supplemental Fig. S1A). In addition, anthocyanin accumulation was also observed in the inflorescence stem and rosette leaves of SAS mutants but not in BAS or MAS mutants (Supplemental Fig. S1, A and B). To investigate the dynamic role of SWI/SNF complexes in regulating chromatin accessibility and gene expression, we performed RNA deep sequencing (RNA-seq) in the inflorescences and subsequently compared the effects of SWI/SNF mutations on gene expression between the seedlings and the inflorescences. The SWI/SNF mutants used in the RNA-seq analysis contain: the BAS mutant *brm*; the SAS mutants *syd*, *swi3d*, and *sys1/2/3*; and the MAS mutants *minu1/2* and *pms2a/b*.

Multi-dimensional scaling (MDS) analyses of the RNA-seq data revealed a high reproducibility of three biological replicates for the inflorescences of each genotype (Fig. 1A). Based on the gene expression patterns, the mutants from the identical SWI/SNF complex are clustered (Fig. 1B), confirming the reliability of the RNA-seq data. We compared differentially expressed genes (DEGs) (|log2FC| ≥1 and FDR < 0.05) detected in the seedlings and inflorescences of SWI/SNF mutants relative to the wild type. Although the SAS mutants had the smaller number of DEGs compared to the BAS and MAS mutants in seedlings (Guo et al., 2022), a significant increase of DEGs was detected in the inflorescences of SAS mutants (Fig. 1C). Except for the weak *syd* mutant allele, the two other SAS mutants *swi3d* and *sys1/2/3* had more DEGs than the BAS and MAS mutants (Fig. 1C; Supplemental Fig. S1C; Supplemental Dataset 1), which is consistent with the fact that the SAS mutants show more severe defects in the flower development than the BAS and MAS mutants (Fig. 1D). This suggests that the SAS complex plays a more important role in inflorescences than in seedlings. We compared DEGs identified in the seedlings and inflorescences of each SWI/SNF mutant and found that only a small portion of DEGs overlap between seedlings and inflorescences (Fig. 1, E-G; Supplemental Fig. S1, D-F), suggesting that the SWI/SNF complexes have a dynamic role in regulating gene expression during development.

**Figure 1.**
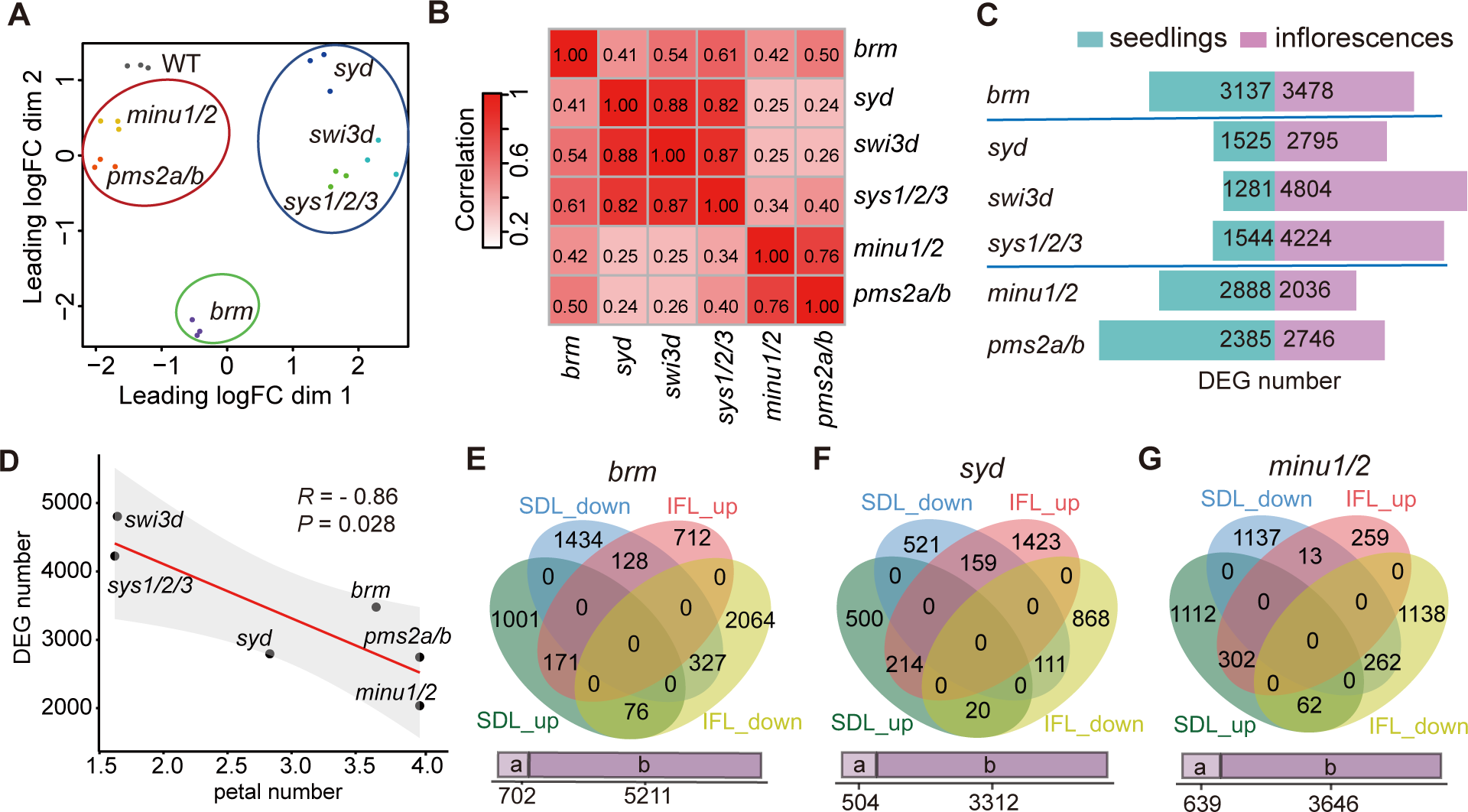
Comparison of DEGs identified by RNA-seq in SWI/SNF mutants between inflorescences and seedlings. **A)** MDS plot of the inflorescence RNA-seq samples. Clustering of BAS, SAS and MAS mutants are marked with green, blue, and red circles, respectively. **B)** Heat map showing the pairwise Pearson correlation coefficients based on the expression fold changes in inflorescences for the total DEGs of the displayed SWI/SNF mutants. **C)** The number of DEGs in SWI/SNF mutants relative to the wild type in both seedlings and inflorescences. **D)** The scatter plot showing the correlation between the average petal number (Guo et al., 2022) and the number of DEGs in inflorescences of the SWI/SNF mutants. A fitted regression line and a 95% confidence interval band are indicated. R represents the Pearson correlation coefficient. *P* value was determined by two-tailed *t* test. **E-G)** The overlap of up-regulated genes and down-regulated DEGs in seedlings and inflorescences of *brm* (**E**), *syd* (**F**) and *minu1/2* (**G**). The purple bars below the venn diagram marked by ‘a’ and ‘b’ represent genes shared by other sets and genes specifically belonging to one set of the venn diagrams, respectively.

### Comparison of SWI/SNF-dependent accessibility between inflorescences and seedlings

Considering that the SWI/SNF complexes are mainly responsible for activating gene expression by promoting chromatin accessibility (Clapier et al., 2017), and given that the three classes of SWI/SNF complexes have been shown to facilitate genome-wide chromatin accessibility in the seedlings (Guo et al., 2022), we determined chromatin accessibility by performing the assay for transposase-accessible chromatin using sequencing (ATAC-seq) and compared the effects of the SWI/SNF mutations on chromatin accessibility between seedlings and inflorescences. On the basis of the principal component analysis (PCA) of ATAC-seq data detected in the inflorescences, the cluster of SAS mutants is more distant from the wild type than the BAS and MAS mutants (Fig. 2A). Consistently, in the inflorescences, down-regulated differentially accessible regions (DARs) identified in the SAS mutants are more abundant than those in the BAS and MAS mutants (Fig. 2B; Supplemental Dataset 2). By comparing the down-regulated differentially accessible regions identified in each SWI/SNF mutant between seedlings and inflorescences, we found that the inflorescence-specific down-regulated differentially accessible regions are also more abundant in the SAS mutants than those in the BAS and the MAS mutants (Fig. 2C). The accessibility changes in inflorescences are strongly correlated among the mutants of the same complex and are weakly correlated among the mutants of different complexes (Fig. 2D). Specially, the fold changes of accessibility in inflorescences are negatively correlated between the SAS mutants and the MAS mutants (Fig. 2D), which is consistent with our previous finding that SAS and MAS have antagonistic effects on regulating chromatin accessibility in seedlings (Guo et al., 2022). These analyses suggest that the three classes of SWI/SNF complexes regulate chromatin accessibility at different genomic regions and that SAS complex contributes more to chromatin accessibility than the other two types of SWI/SNF complexes in inflorescences.

**Figure 2.**
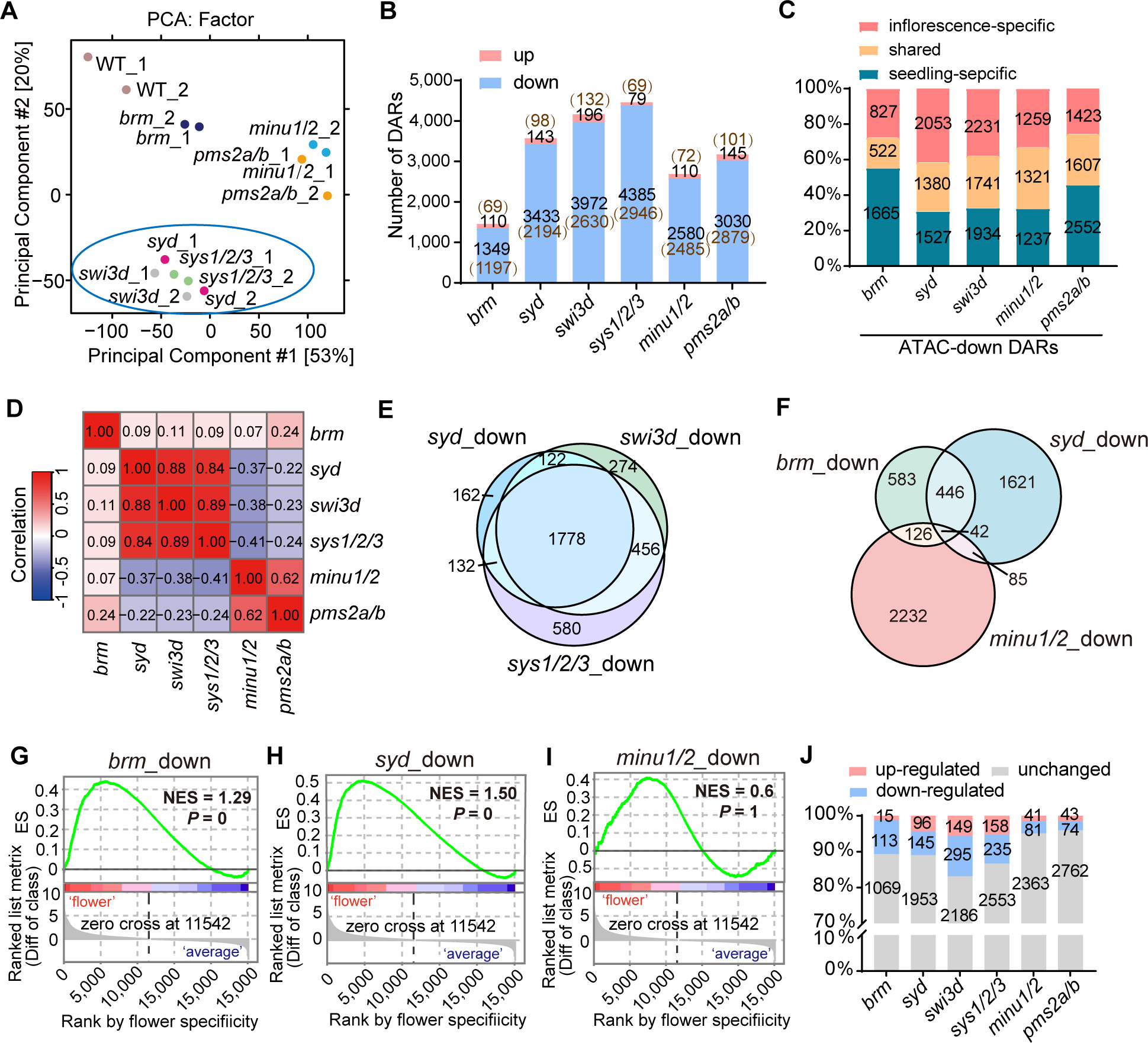
Analysis of differentially accessible regions in inflorescences of SWI/SNF mutants relative to the wild type. **A)** Principal component analysis (PCA) of the inflorescence ATAC-seq data of the wild type and SWI/SNF mutants. The cluster of SAS mutants are marked by the blue circle, respectively. **B)** The number of up-regulated and down-regulated differentially accessible regions in inflorescences of the SWI/SNF mutants identified by ATAC-seq. The number of genes annotated by the differentially accessible regions is indicated in brackets with brown font. DARs, differentially accessible regions. **C)** The proportion and number of inflorescent-specific, seedling-specific and shared differentially accessible regions among total differentially accessible regions identified in the inflorescences and seedlings of each SWI/SNF mutant compared to the wild type. DARs, differentially accessible regions. **D)** Heat map showing the pairwise Pearson correlation coefficient based on the accessibility changes at the total genes with changed accessibility in inflorescences of the displayed SWI/SNF mutants. **E)** Venn diagram displaying the overlap of genes with down-regulated accessibility in inflorescences of SAS mutants *syd*, *swi3d* and *sys1/2/3* compared to the wild type. **F)** Venn diagram showing the overlap of genes with down-regulated accessibility in inflorescences of *brm*, *syd* and *minu1/2* compared to the wild type. **G-I)** Line plots depicting the enrichment of flower-specific expressed genes among the genes with down-regulated accessibility in *brm* (**G**), *syd* (**H**) and *minu1/2* (**I**) as determined by GSEA analysis. The grey regions represent the ranked list of the differences between the expression in flowers and the average expression in all tissues of each Arabidopsis gene. The normalized enrichment scores were labeled. *P* values were determined by permutation. **J)** The proportion of genes showing up-regulated, down-regulated, and unchanged expression among genes with down-regulated accessibility in the inflorescences of the indicated SWI/SNF mutants relative to the wild type.

Furthermore, we analyzed the distribution of the down-regulated differentially accessible regions in both seedlings and inflorescences of each SWI/SNF mutant and found that the distributions are overall identical between the seedlings and inflorescences (Supplemental Fig. S2, A and B; Supplemental Dataset 2). While the BAS and MAS-regulated accessible regions are enriched at the TSS-flanking regions, the SAS-regulated accessible regions are enriched at distal promoter regions and intergenic regions, which we hereafter refer to as distal accessible regions (Supplemental Fig. S2, A and B). In both seedlings and inflorescences, the SAS complex facilitates the accessibility at ∼50% of distal accessible regions, whereas BAS and MAS mediate the accessibility only at < 10% of them (Supplemental Fig. S2C). In particular, the SAS complex contributes to chromatin accessibility at ∼40% of the super enhancer regions defined by the top 2.5% of longest nonpromotor accessible region clusters in a previous study in Arabidopsis (Zhao et al., 2022), whereas the BAS and MAS complexes only show a marginal effect on the accessibility of the super enhancer regions (Supplemental Fig. S2D). These results point to the important role of SAS in promoting chromatin accessibility at distal accessible regions in both seedlings and inflorescences.

The ATAC-seq analysis indicated that the genes with decreased accessibilities are highly overlapped in different SAS mutants (Fig. 2E), suggesting the SAS components function together in regulating chromatin accessibility. Although the BAS and SAS complexes regulate chromatin accessibility primarily at different genomic regions (Fig. 2D; Supplemental Fig. S2, A and B), a substantial subset (∼40%) of genes with down-regulated accessibility in the BAS mutant are shared by those with down-regulated accessibility in the SAS mutant (Fig. 2F). Given that the BAS and SAS can regulate the accessibility of the same set of target genes at proximal and distal regions, respectively (Supplemental Fig. S2, A-C), the BAS and SAS complexes are likely to collaborate in regulating the accessibility of different regions at a subset of their shared target genes.

### SAS regulates the accessibility and transcription of genes involved in flower development

We conducted a gene set enrichment analysis (GSEA) to examine genes with down-regulated accessibility in the inflorescences of the BAS, SAS, and MAS mutants. We found that flower-specific expressed genes exhibit enrichment among the genes with down-regulated accessibility in the SAS mutant, and to a lesser extent in the BAS mutant, whereas we did not observe this enrichment in the MAS mutant (Fig. 2, G-I; Supplemental Fig. S3A). Consistently, the Gene Ontology (GO) analysis revealed a significant enrichment of flower development-related genes among those exhibiting down-regulated accessibility in the BAS and SAS mutants, and such enrichment was not observed in the MAS mutants (Supplemental Fig. S3A; Supplemental Dataset 3). Interestingly, the genes with down-regulated accessibility in the MAS mutants showed enrichment in terms associated with protein targeting and metabolic processes (Supplemental Fig. S3A; Supplemental Dataset 3). The housekeeping genes identified by the previous study (Cheng et al., 2017) are significantly more enriched among genes showing down-regulated accessibility in the MAS mutant than in the BAS and SAS mutants (Supplemental Fig. S3B), suggesting that MAS shows the tendency to regulate housekeeping genes but BAS and SAS avoid to regulate these genes. These results indicate that SAS and BAS complexes but not the MAS complex are responsible for promoting the accessibility of genes involved in flower development.

Among the genes with down-regulated accessibility in the inflorescences of each SWI/SNF mutant, the DEGs are most abundant in the SAS mutants, followed by the BAS mutant and then the MAS mutants (Fig. 2J). Moreover, the DEGs with decreased expression were more abundant than those with increased expression in each SWI/SNF mutant (Fig. 2J), supporting the notion that SWI/SNF-dependent accessibility primarily functions in activating transcription rather than repressing it. It is noteworthy that most of the genes with down-regulated accessibility in SWI/SNF mutants did not show changes in expression (Fig. 2J), indicating that the SWI/SNF-dependent accessibility contributes to establishing a priming chromatin state for transcription but is not sufficient for actual transcription to occur. Taken together, these results suggest that among the three classes of SWI/SNF complexes, the SAS complex plays a major role in promoting chromatin accessibility and facilitating transcriptional activation during flower development.

### SAS is responsible for establishing the accessibility of genes involved in flower development

The dynamic regulation of chromatin accessibility is involved in the regulation of gene expression during flower development (Yan et al., 2019). To comprehensively investigate the accessibility of genes involved in flower development at the whole-genome level, we compared the chromatin accessibility between seedlings and inflorescences based on our ATAC-seq results in the wild type. The ATAC-seq analysis identified a total of 1,325 genes (corresponding to 1,842 peaks) with increased accessibility and 1,344 genes (corresponding to 1,742 peaks) with decreased accessibility in inflorescences compared to seedlings (Fig. 3A, Supplemental Dataset 4). The genes displaying increased accessibility in inflorescences were found to be enriched with gene ontology (GO) terms associated with flower development (Supplemental Fig. S4A). Conversely, the genes exhibiting decreased accessibility were enriched with GO terms related to stress response (Supplemental Fig. S4B).

**Figure 3.**
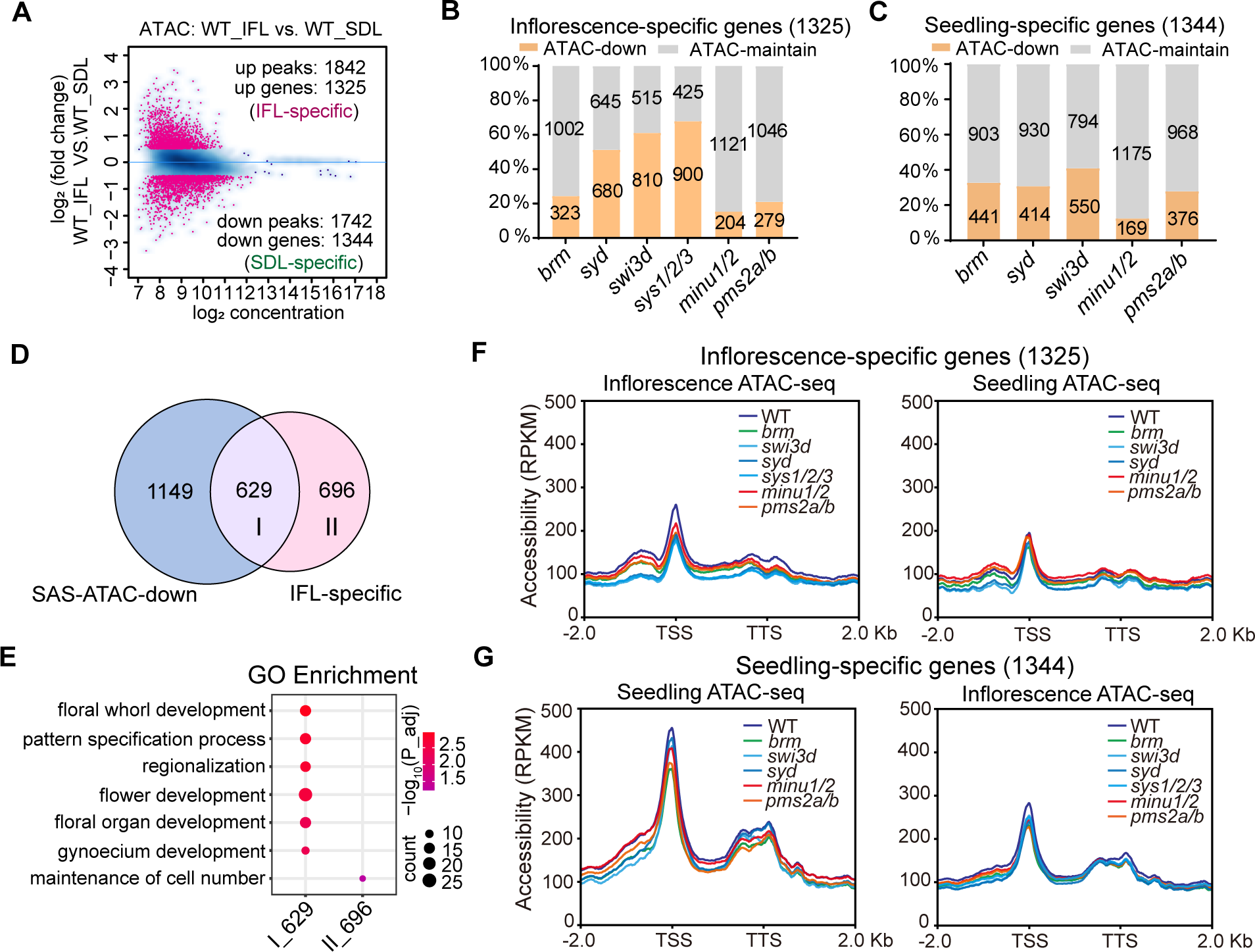
Effect of BAS, SAS, and MAS mutations on the accessibility of seedling-specific and inflorescence-specific accessible regions. **A)** M (log ratio) versus A (log average) (MA) plot showing the differential analysis results of ATAC-seq peaks in inflorescences versus in seedlings of the wild-type plants. The plot shows the log_2_ (fold change) and log_2_ (average reads concentration) of the ATAC-seq peaks. The red points represent the significant changed peaks with FDR<0.05 and |log_2_ (fold change) | ≥ 0.5. **B)** The percentage of genes with down-regulated accessibility in the inflorescences of indicated SWI/SNF mutants among the inflorescence-specific accessible genes. **C)** The percentage of genes with down-regulated accessibility in the seedlings of indicated SWI/SNF mutants among the seedling-specific accessible genes. **D)** Venn diagram showing the overlap of the genes with consistently down-regulated accessibility in the inflorescences of SAS mutants (*syd*, *swi3d*, and *sys1/2/3*) compared to the wild type and the inflorescence-specific accessible genes. Gene set I represents the inflorescence-specific accessible genes regulated by SAS, and gene set II represents the inflorescence-specific accessible genes not regulated by SAS. **E)** Bubble plot showing the enriched GO terms for gene set I and gene set II defined in (**D**). The size of the bubble represents the gene number, and the color of the bubble represents the adjusted *P* value of the enriched term. **F, G)** Profile plots depicting the ATAC-seq signals of the SWI/SNF mutants and the wild-type plants in inflorescences and in seedlings at the genic regions with inflorescence-specific accessibility (**F**) and the genic regions with seedling-specific accessibility (**G**).

Next, we determined to what extent the SWI/SNF complexes contribute to the accessibility of inflorescence-specific accessible genes, and found that > 50% of the inflorescence-specific accessible genes were regulated by the SAS components, while around 20% were regulated by the BAS or MAS components (Fig. 3B). In contrast, the numbers of seedling-specific accessible genes regulated by BAS, SAS, and MAS were not markedly different (Fig. 3C). Among the 1,325 inflorescence-specific accessible genes, 47% (629/1,325) of them are overlapped with the 1,778 genes with decreased accessibility in all the three tested SAS mutants (Fig. 3D). The GO analysis indicated that the 629 SAS-regulated, inflorescence-specific accessible genes (type I) are closely associated with flower development, whereas the remaining 696 inflorescence-specific accessible genes (type II) not regulated by SAS were enriched with the term ‘maintenance of cell number’, which is not directly related to flower development (Fig. 3E), confirming that SAS is specifically responsible for establishing flower-specific chromatin accessibility.

To characterize the accessible regions regulated by three classes of SWI/SNF complexes in both seedlings and inflorescences, we drew the average accessible signals at the 1,325 inflorescence-specific accessible genes and the 1,344 seedling-specific accessible genes in both wild-type plants and SWI/SNF mutants. For the inflorescence-specific accessible genes, we observed not only a TSS-flanking accessible region but also a distal accessible region upstream of the TSS (Fig. 3F, Supplemental Fig. S5A). This characteristic aligns with the typical feature of SAS-regulated accessible genes (Guo et al., 2022). However, the seedling-specific accessible genes do not exhibit this distinct ‘SAS feature’ (Fig. 3G; Supplemental Fig. S5B). SAS is the primary regulator required for the accessibility of inflorescence-specific accessible genes, whereas BAS, SAS, and MAS collectively contribute to the accessibility of seedling-specific genes (Fig. 3G; Supplemental Fig. S5B). It is worth noting that SAS weakly contributes to the accessibility of inflorescence-specific accessible genes in seedlings (Fig. 3F; Supplemental Fig. S5A). However, in wild-type plants, the accessibility of these genes is markedly lower in seedlings than in inflorescences (Fig. 3F; Supplemental Fig. S5A). Therefore, SAS contributes to chromatin accessibility at the distal promoter region of genes related to flower development.

### Comparison of the binding patterns of BAS, SAS, and MAS in inflorescences and seedlings

To investigate the molecular mechanism underlying the different effects of SWI/SNF complexes on chromatin accessibility between seedlings and inflorescences, we conducted chromatin immunoprecipitation followed by sequencing (ChIP-seq) for the BAS subunit SWI3C, the SAS subunit SWI3D, and the MAS subunit PMS2B. We observed that the replicates of the same sample are more correlated than different samples (Supplemental Fig. S6A), indicating the reliability of our ChIP-seq results. The results indicated that the binding peaks of the BAS and SAS subunits in both seedlings and inflorescences formed a cluster with strong correlation, while displaying weak correlation with the binding peaks of the MAS subunit (Fig. 4A). However, upon closer examination, we found that the binding peaks of BAS and SAS subunits within the same developmental stage exhibited higher correlation than one subunit across different developmental stages (Fig. 4A), suggesting that BAS and SAS exhibited similar distribution patterns in either seedlings or inflorescences, and their distributions showed differences between the two developmental stages. Analysis of the binding peaks called from the ChIP-seq data revealed that the three SWI/SNF complexes displayed a similar number of binding peaks between seedlings and inflorescences (Fig. 4B; Supplemental Dataset 5). Previous studies have indicated that, in seedlings, BAS and SAS share the majority of their binding peaks, while most of the binding peaks of MAS are different from those of BAS and SAS (Guo et al., 2022; Fu et al., 2023). The overlapping patterns of BAS, SAS, and MAS peaks in inflorescences were similar to those in seedlings (Supplemental Fig. S6B). However, whether and how the binding of SWI/SNF complexes to chromatin is altered during plant development is largely unclear.

**Figure 4.**
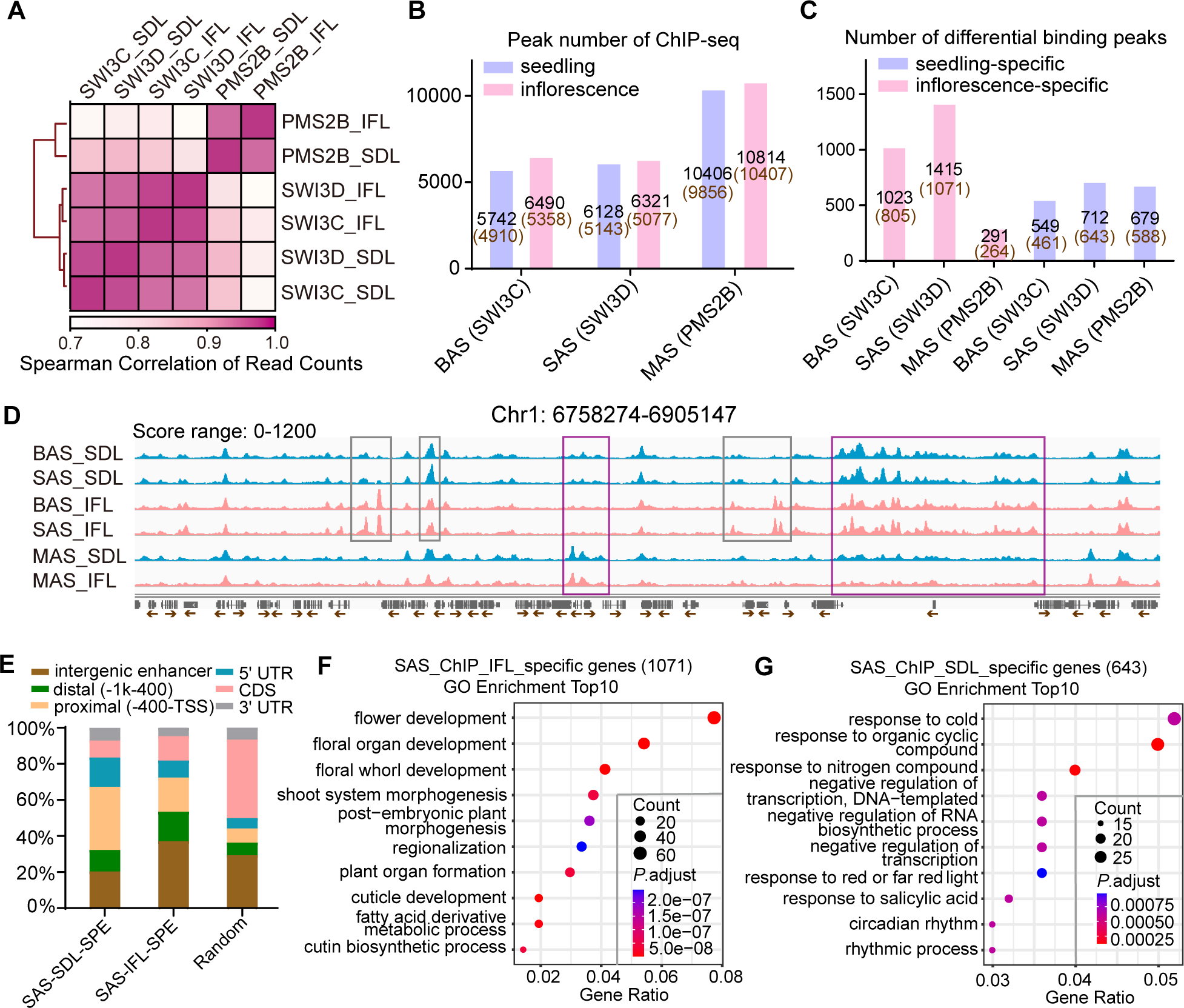
Specifically binding regions of BAS, SAS, and MAS in seedlings and inflorescences. **A)** Heat map showing the pairwise Spearman correlation coefficients based on the ChIP-seq read counts of the representative subunit of BAS (SWI3C), SAS (SWI3D), and MAS (PMS2B) in seedlings (SDL) and inflorescences (IFL). **B)** Peak number determined by ChIP-seq for the subunits of BAS, SAS and MAS in seedlings and inflorescences. The number of genes annotated by the peaks is provided in brackets with brown font. **C)** Number of seedling-specific and inflorescence-specific binding peaks for the BAS, SAS and MAS subunits. The number of genes annotated by the peaks is provided in brackets with brown font. **D)** Snapshot showing the ChIP-seq signals of BAS, SAS and MAS subunits in inflorescences and seedlings at a genomic region. Blue peaks represent ChIP-seq signal in seedlings, and pink peaks represent ChIP-seq signal in inflorescences. Peaks marked with grey boxes indicate differential binding regions of BAS/SAS subunits between seedlings and inflorescences. Peaks marked with purple boxes indicate differential binding regions between the MAS subunit and the BAS/SAS subunits. The scale of normalized reads is shown. **E)** The proportion of regions annotated to specified chromatin features. The left two bars represent regions specifically bound by SAS in seedlings and inflorescences. Random: 8,000 randomly selected genomic sites. **F, G)** Bubble plots showing the top 10 enriched GO terms in genes specifically bound by SAS in inflorescences (**F**) and in seedlings (**G**).

By comparing the binding peaks of each SWI/SNF subunit between seedlings and inflorescences, we identified 1,023 BAS peaks, 1,415 SAS peaks, and 291 MAS peaks exhibiting increased accumulation in inflorescences relative to seedlings, which we termed inflorescence-specific binding peaks; and revealed 549 BAS peaks, 712 SAS peaks, and 679 MAS peaks with decreased accumulation in inflorescences compared to seedlings, which we termed seedling-specific binding peaks (Fig. 4C; Supplemental Fig. S6, C-E; Supplemental Dataset 6). Although the three SWI/SNF complexes showed similar numbers of seedling-specific binding peaks, the number of inflorescence-specific binding peaks was significantly higher for SAS and BAS compared to MAS (Fig. 4B). This suggests that the binding of SAS and BAS to chromatin undergoes dynamic establishment during flower development. We assigned the binding peaks to nearby genes and then defined the binding genes for the three SWI/SNF complexes (Fig. 4, B and C), and found that the BAS and SAS-binding genes were significantly overlapped among either inflorescence- or seedlings-specific binding genes, whereas only a small portion of MAS-binding genes were overlapped with BAS or SAS-binding genes (Supplemental Fig. 6, F and G). This was exemplified by the binding signals of BAS, SAS, and MAS in seedlings and inflorescences at their random target genomic regions (Fig. 4D). These results suggest that the binding of BAS and SAS to chromatin is altered during development at a substantial subset of their target loci.

Furthermore, we determined whether the distribution of the binding peaks for each SWI/SNF complex across the genic region is altered in inflorescences compared to that in seedlings, and found that the overall distribution of total binding peaks for each SWI/SNF complex is similar between inflorescences and seedlings (Supplemental Fig. S6H). However, the proportion of the SAS and BAS-binding peaks in intergenic and distal promoter regions exhibited a slight increase in inflorescences relative to seedlings (Supplemental Fig. S6H). Upon comparing the distributions of inflorescence-specific SAS-binding regions and seedling-specific SAS-binding regions, we observed that SAS has a higher tendency to bind intergenic and distal promoter regions in inflorescences than in seedlings (Fig. 4E). The GO analysis revealed that the inflorescence-specific SAS-binding genes are primarily associated with flower development, while the seedling-specific SAS-binding genes are enriched in terms of stress response (Fig. 4, F and G). These results suggest that the SAS is specifically recruited to a subset of genes involved in flower development during the reproductive phase, rather than maintaining a constant association with these genes between the vegetative and reproductive processes.

### Recruitment of SAS establishes the accessibility of flower-development genes

To determine the impact of BAS, SAS, and MAS binding on gene accessibility in inflorescences and seedlings, we examined the genes specifically bound by each complex in either inflorescences or seedlings and compared them to genes that exhibited decreased accessibility in the corresponding complex mutant. Interestingly, we found that inflorescence-specific SAS-binding genes showed the highest proportion of accessibility changes compared to the inflorescence-specific BAS- and MAS-binding genes (Fig. 5A). It is worth noting that although the numbers of inflorescence-specific binding genes were comparable for BAS and SAS, the proportion of genes with down-regulated accessibility was significantly lower in the BAS mutant (95/805) than in the SAS mutant (382/1,071). This discrepancy in accessibility changes between the BAS and SAS mutants might be attributed to the different remodeling activities of the two complexes at their target regions. Further examination of SAS-specific binding regions in inflorescences revealed that the regions with decreased accessibility exhibited higher binding and accessibility signals compared to regions with maintained accessibility (Fig. 5, B and C). This suggests that the enhanced binding affinity of SAS at specific binding regions in inflorescences promotes the remodeling activity, leading to increased accessibility.

**Figure 5.**
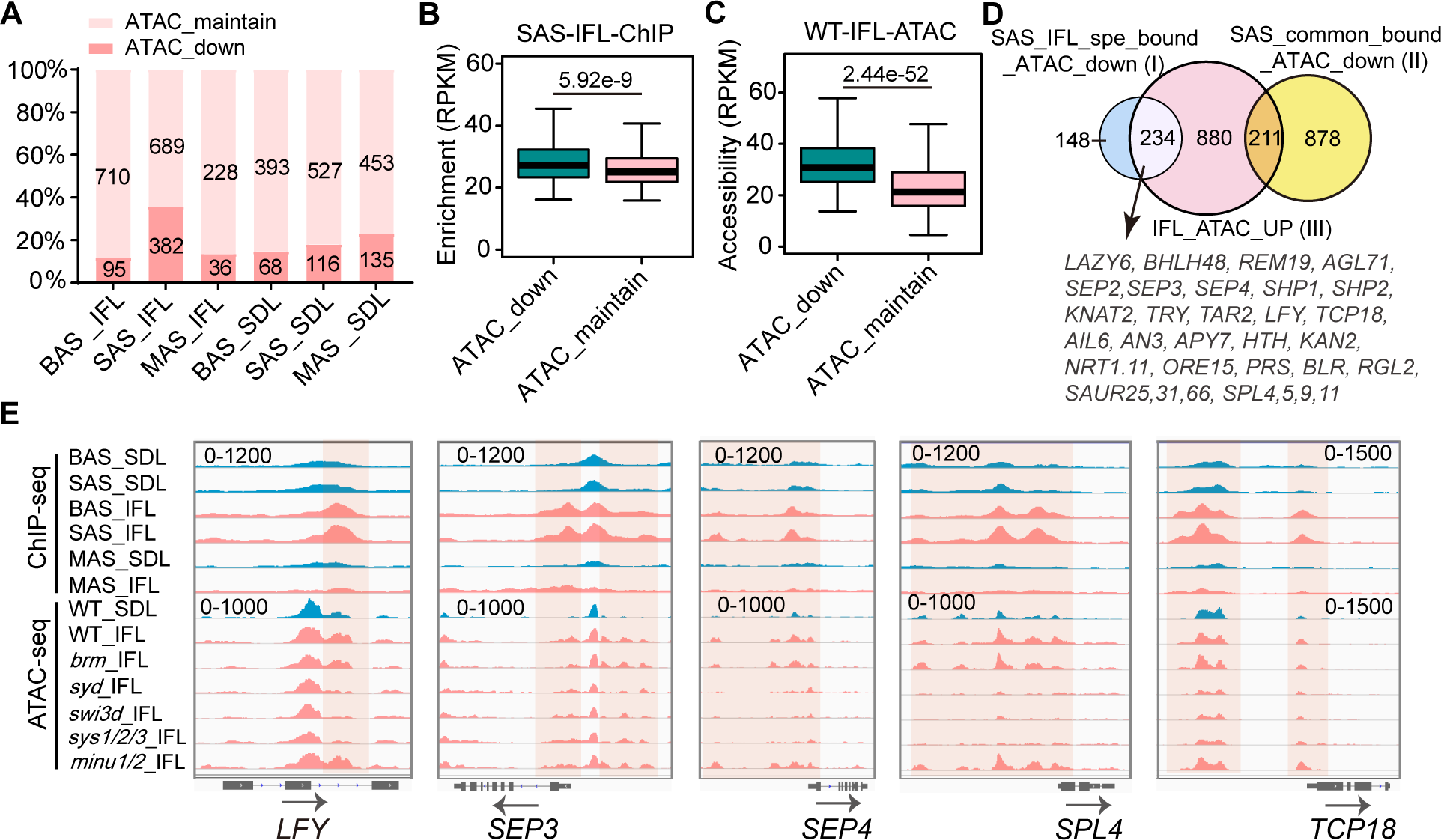
Specific binding of SAS in inflorescence contributes to establishing the accessibility of flower development-related genes. **A)** The proportion of genes with down-regulated accessibility and maintained accessibility in the corresponding mutants among genes specifically bound by BAS, SAS and MAS in inflorescences (IFL) and in seedlings (SDL). **B, C)** Boxplots showing the enrichment of SAS in inflorescences (**B**) and the accessibility in the inflorescences of the wild-type plants (**C**) at the regions specifically bound by SAS in inflorescences with down-regulated accessibility (green) or maintained accessibility (pink) in the SAS mutant. The center lines and box edges of the box plots represents medians and the interquartile range (IQR), respectively. Whiskers extent within 1.5 times the IQR. *P* values were determined by two-tailed Mann-Whitney *U* test. **D)** Venn diagram showing the overlap between three gene sets: (I) genes specifically bound by SAS in inflorescences with decreased accessibility in the SAS mutant, (II) genes bound by SAS in both seedlings and inflorescences with decreased accessibility in the SAS mutant, and (III) the inflorescence-specific accessible genes. Devel-opment-related genes selected from the overlap between gene sets I and III are listed below the Venn diagram. **E)** Snapshots displaying the ChIP-seq signal of the three SWI/SNF complexes and the ATAC-seq signal of SWI/SNF mutants in inflorescences and in seedlings at the genic region of genes involved in flower development. Blue represents the signal in seedlings, and pink represents the signal in inflorescences. Differential peaks of ChIP-seq and ATAC-seq in seedlings and inflorescences are marked by shadows. The scale of normalized reads is shown in each snapshot.

To identify inflorescence-specific accessible genes that are specifically bound and remodeled by SAS, we conducted an overlap analysis between the 382 SAS-specific binding genes with decreased accessibility in the SAS mutant inflorescences (Fig. 5A) and the 1,325 inflorescence-specific accessible genes (Fig. 3A). Remarkably, we found that 61% (234/382) of the genes specifically bound and remodeled by SAS in inflorescences were inflorescence-specific accessible genes, accounting for 18% (234/1,325) of the total inflorescence-specific accessible genes (Fig. 5D). This subset includes crucial flower development-related genes, such as *LFY* (Schultz and Haughn, 1991), *SEP2/3/4* (Pelaz et al., 2000; Ditta et al., 2004), *SAUR*s (Stortenbeker and Bemer, 2019), and *SPL*s (Chen et al., 2010) (Fig. 5D). Snapshots illustrating the association between SWI/SNF binding and SWI/SNF-dependent accessibility at representative genes are provided (Fig. 5E). These analyses highlight the contribution of SAS recruitment to the establishment of accessibility for flower development-related genes during flower development. Additionally, we discovered that 16% (211/1,325) of the inflorescence-specific accessible genes exhibited SAS-dependent accessibility but were bound by SAS in both inflorescences and seedlings (Fig. 5E), suggesting that the involvement of SAS in inflorescence-specific accessibility is also regulated after SAS is recruited to its target genes.

Furthermore, we investigated whether the SAS complex is involved in the expression of flower development-related genes. We found that the accessibility of inflorescence-specific accessible regions is significantly lower in the SAS mutant than in the wild-type in the inflorescences (Supplemental Fig. S7A), confirming that the SAS complex is crucial for facilitating the accessibility of the flower development-related genes specifically in inflorescences. As determined by our RNA-seq data, the expression levels of inflorescence-specific accessible genes are significantly higher in inflorescences than in seedlings, and their expression levels are reduced in the SAS mutant compared to the wild-type in inflorescences (Supplemental Fig. S7B). In the SAS mutant, the inflorescence-specific accessible genes with decreased expression are more abundant than those with increased expression (Supplemental Fig. S7C). These results suggest that the SAS complex contributes to chromatin accessibility and thereby facilitates the transcription of flower development-related genes. However, we also found that the expression levels of a substantial subset of inflorescence-specific accessible genes are not significantly reduced in the SAS mutant (Supplemental Fig. S7C). Therefore, the SAS-dependent chromatin accessibility observed in inflorescences may result in transcription occurring exclusively in specific cells and developmental stages, and these localized transcriptional events may not be detectable in the RNA-seq analysis of mixed inflorescence tissues.

### SAS-dependent accessible regions and transcription factors

Similar to seedlings, where most SWI/SNF-regulated accessible regions are direct targets of SWI/SNF complexes (Guo et al., 2022), we observed that in the inflorescences of SWI/SNF mutants, the majority of accessible regions with decreased accessibility are also directly bound by their corresponding SWI/SNF complexes (Fig. 6A). Out of the 3,972 accessible regions with decreased accessibility detected in *swi3d* inflorescences, 60% of them are bound by SAS. Specifically, 14% of these regions are exclusively bound by SAS in inflorescences, while 46% are bound by SAS in both seedlings and inflorescences (Fig. 6B). Comparison of the inflorescence-specific SAS-binding regions with the shared SAS-binding regions in seedlings and inflorescences revealed that the inflorescence-specific regions exhibit higher SAS-binding signals and lower accessibility signals (Fig. 6, C and D). It is important to note that the inflorescence-specific binding of SAS to its target loci only accounts for a small subset of SAS-dependent accessible regions (Fig. 6B). This further supports the notion that the regulation of SAS-dependent accessibility in inflorescences involves additional downstream steps following the binding of SAS to its target loci.

**Figure 6.**
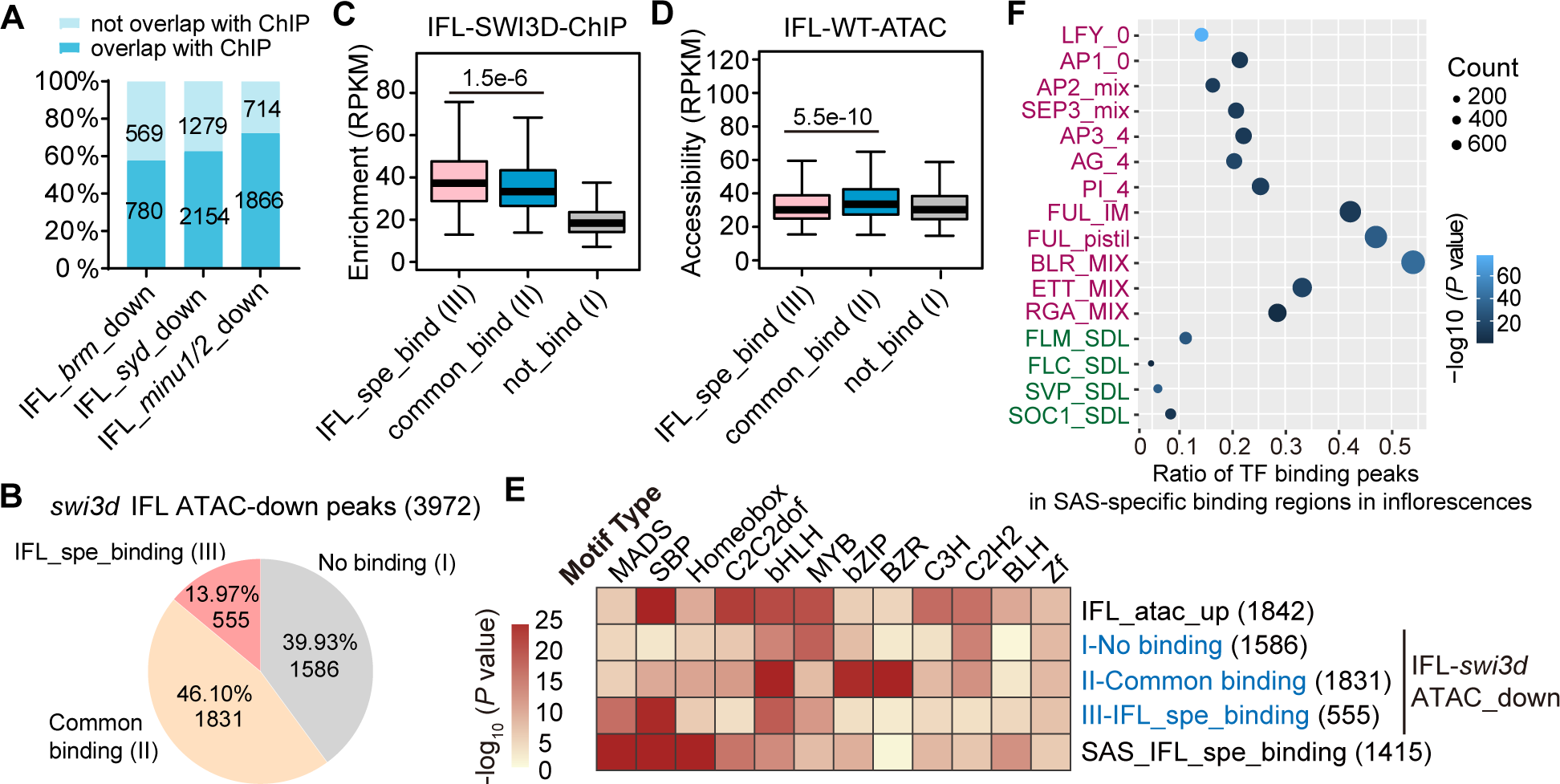
Characterization of SAS-dependent accessible regions in inflorescences. **A)** The proportion of the BAS/SAS/MAS bound and unbound regions among regions with down-regulated accessibility in inflorescences of the core enzyme mutants of BAS, SAS and MAS. **B)** Pie chart showing the proportion of the peaks not bound by SAS (I), commonly bound by SAS in both seedlings and inflorescences (II) and specifically bound by SAS in inflorescences (III) among the total down-regulated peaks in the inflo-rescences of swi3d relative to the wild type. **C, D)** Boxplots showing the enrichment of SWI3D (**C**) and the accessibility in the inflorescences of the wild-type plants (**D**) at the regions I, II and III defined in Figure 5F. The center lines and box edges of the box plots represents medians and the interquartile range (IQR), respectively. Whiskers extend within 1.5 times the IQR. P values were determined by two-tailed Mann-Whitney *U* test. **E)** Heatmap exhibiting the different enrichment of the different enrichment of various motifs at the indicated regions. **F)** Bubble plot showing the ratio of the transcription factor (TF) binding peaks collected by the previous studies (Chen et al., 2018; van Mourik et al., 2023) among the 1,415 specific binding peaks of SWI3D. The binding peaks of TFs in inflorescences and seedlings are marked in purple and green, respectively.

Previous studies have demonstrated that transcription factors either recruit SWI/SNF complexes to specific target genes or collaborate with SWI/SNF to regulate chromatin accessibility and transcriptional activation (Bieluszewski et al., 2023). In order to investigate how SAS-dependent chromatin accessibility is regulated during flower development, we examined whether the inflorescence-specific SAS-binding regions are enriched with any transcription factor-binding DNA motifs. Interestingly, we found that the inflorescence-specific SAS-binding regions are significantly enriched with the DNA motifs bound by MADS and SBP transcription factors (Fig. 6E; Supplemental Dataset 7). Given that the MADS and SBP transcription factors are important for flower development (Chen et al., 2010; Theißen et al., 2016), it is likely that these transcription factors either facilitate the recruitment of SAS to flower development-related genes or collaborate with SAS to regulate the accessibility and/or transcription of these genes. To further explore the relationship between SAS and the MADS-family transcription factors, we compared the inflorescence-specific SAS-binding regions with the MADS-binding regions previously identified in inflorescences or seedlings (Chen et al., 2018; van Mourik et al., 2023). The analysis indicated that the inflorescence-specific SAS-binding regions exhibited a high level of overlap (15%∼50%) with the regions bound by multiple MADS transcription factors in inflorescences, whereas the overlap (<10%) was markedly reduced when they are compared with the regions bound by the MADS transcription factors detected in seedlings (Fig. 6F). These findings support the idea that MADS transcription factors collaborate with the SAS complex to regulate the transcription of genes involved in flower development.

### The accessibility generated by SAS in inflorescence is required for AP1 binding

We chose AP1, one famous MADS transcription factor crucial for floral identity, as an example to study the relationship between MADS transcription factors and SAS in inflorescences. At the genomic regions showing down-regulated accessibility in the SAS mutants, the ATAC-seq signals in the *ap1* mutant are overall not affected compared to the wild-type plants in inflorescences (Fig. 7A). There are few differential accessible regions in *ap1* compared with the wild-type plants in inflorescences (Fig. 7B). These data indicate that AP1 is unlikely to function upstream to the SAS complex.

**Figure 7.**
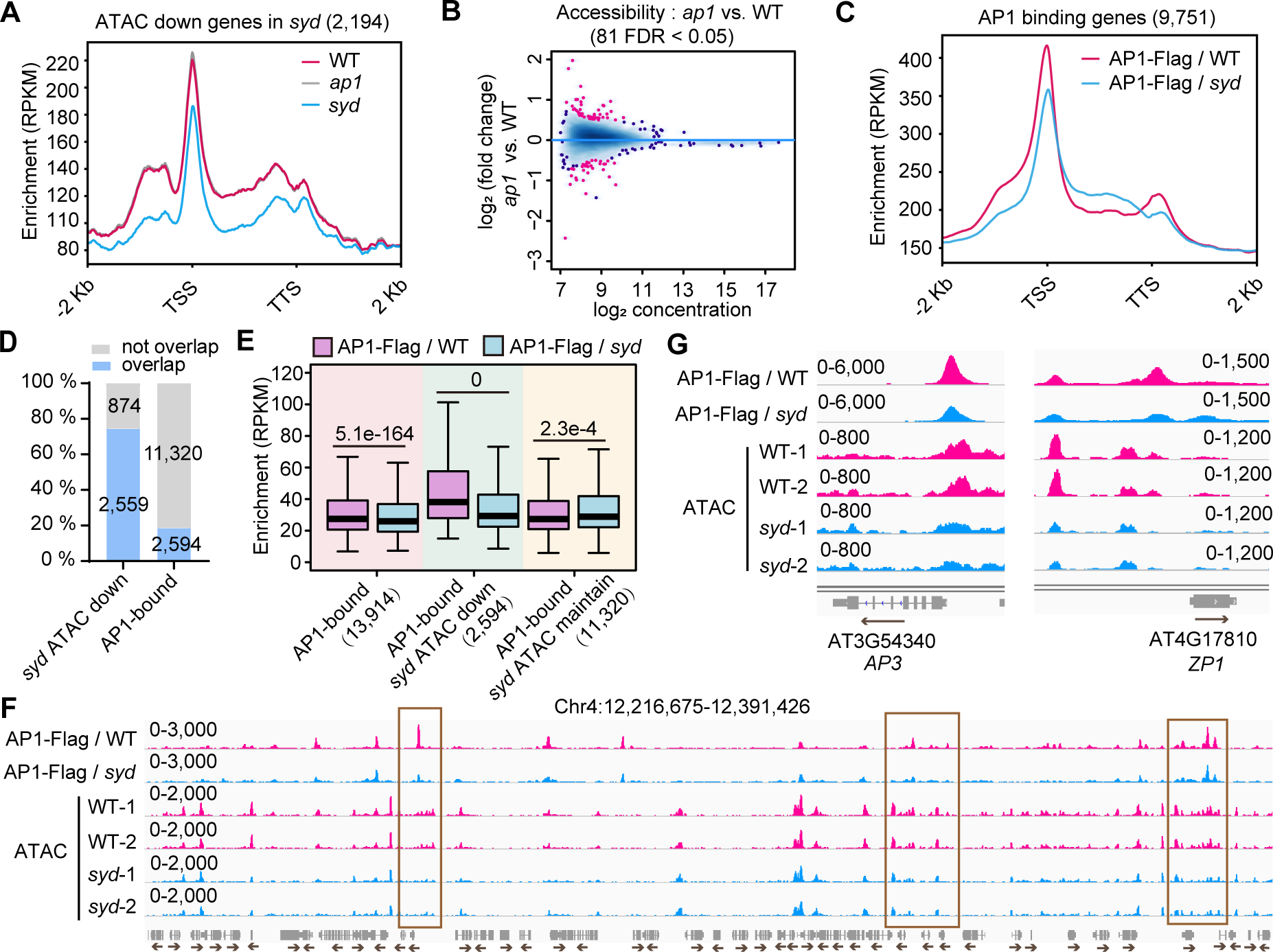
The binding of AP1 depends on the accessibility generated by SAS in inflorescences. **A)** Profile plots depicting the ATAC-seq signals in inflorescences of *ap1*, *syd* and the wild-type plants at the genic regions with down-regulated accessibility in *syd*. **B)** MA plot showing the differential analysis results of ATAC-seq peaks in inflorescences of *ap1* versus the wild-type plants. The plot shows the log_2_ (fold change) and log_2_ (average reads concentration) of the ATAC-seq peaks. The red points represent the significant changed peaks with FDR<0.05. **C)** Profile plots depicting the ChIP-seq signals of AP1-Flag in the background of wild-type and the *syd* mutant at the genic regions bound by AP1 in inflorescences. **D)** The percentage of the down-regulated accessible regions in *syd* overlapped with the AP1-bound regions and the percentage of the AP1-bound regions overlapped with the down-regulated accessible regions in *syd.* The number of the regions are marked on the bar plots. **E)** Boxplots showing the enrichment of AP1 in the wild type and the *syd* mutants at the AP1-bound regions, AP1-bound regions with down-regulated accessibility in *syd* and AP1-bound regions with maintained accessibility in *syd*, respectively. The center lines and box edges of the box plots represents medians and the interquartile range (IQR), respectively. Whiskers extent within 1.5 times the IQR. *P* values were determined by two-tailed Wilcoxon signed rank test. **F)** Snapshot demonstrating the ChIP-seq and ATAC-seq signals of the indicated samples at a genomic region. The down-regulated binding peaks of AP1 at the regions with down-regulated accessibility in SAS mutants are marked by the brown boxes. **G)** Snapshot demonstrating the ChIP-seq and ATAC-seq signals of the indicated samples at two genes targeted by AP1. In (**F**) and (**G**), the scale of normalized reads is shown.

We therefore investigated whether AP1 functions downstream of SAS by performing ChIP-seq of AP1 in the inflorescences of wild-type and *syd* mutant plants. The ChIP-seq data indicated that the average enrichment of AP1 is markedly decreased in *syd* compared to the wild type (Fig. 7C), suggesting that the genomic binding of AP1 is partially dependent on SAS. By overlapping the AP1-binding regions and the regions showing down-regulated accessibility in the *syd* mutant, we found most of the accessible regions regulated by SAS in inflorescences are bound by AP1 (Fig. 7D). Among the binding regions of AP1, the regions with down-regulated accessibility in *syd* show a significant higher AP1 enrichment than the regions with maintained accessibility (Fig. 7E), and the enrichment of AP1 in SAS-regulated accessible regions is significantly decreased in the *syd* mutant compared to the wild type (Fig. 7E). The effects of *syd* on chromatin accessibility and the genomic binding of AP1 were exemplified by the screenshot of a representative genomic region (Fig. 7F). In addition, *AP3* (Hill et al., 1998) and *ZP1* (Hu et al., 2023), two well-known target genes of AP1, show consistent decrease of chromatin accessibility and AP1 enrichment in the *syd* mutant relative to the wild type (Fig.7G). These data suggest that the SAS-dependent chromatin accessibility contributes to the binding of AP1 to its target genomic loci.

## Discussion

SWI/SNF complexes play crucial roles in the regulation of diverse developmental processes in both metazoans and plants. Recent studies have illustrated the functional variations among the three Arabidopsis SWI/SNF complexes: BAS, SAS, and MAS (Guo et al., 2022; Fu et al., 2023). However, the precise mechanisms by which these complexes exert specific functions during different developmental stages remain largely unknown. In this study, we performed a comprehensive analysis to compare the impact of BAS, SAS, and MAS on chromatin accessibility throughout the entire genome between vegetative and reproductive development processes. Our results demonstrate that while the three SWI/SNF complexes contribute to a comparable amount of accessible regions in seedlings, there is a significantly higher proportion of inflorescence-specific accessible regions that depend on SAS. This indicates that SAS exhibits a stronger influence on inflorescences compared to BAS and MAS. The findings highlight the specific role of SAS in inflorescences, which is consistent with the more severe defects in floral development observed in SAS mutants relative to BAS and MAS mutants.

Our analysis, combining ATAC-seq with ChIP-seq data, implicates two mechanisms through which the SAS complex contributes to establishing the accessibility of genes involved in flower development. Firstly, SAS is recruited to genes involved in flower development, thereby establishing the accessibility of these genes. It is likely that multiple transcription factors collaborate to recruit SAS to different sets of flower development related genes. Secondly, the SAS complex exhibits constant binding to genes involved in flower development during both the vegetative and reproductive processes, with an enhanced chromatin remodeling activity specifically during the reproductive process. The change in the chromatin remodeling activity of SAS at different developmental stages can be attributed to either its differential recognition ability to histone modifications or its interactions with different transcription factors. Further studies are needed to investigate how histone modifications and/or transcription factors regulate the recruitment of SAS to chromatin or its chromatin remodeling activity during the reproductive process.

A previous study have indicated that establishing the accessibility of distal accessible regions is critical for the activation of genes involved in flower development during reproductive development (Yan et al., 2019). However, the specific mechanisms responsible for establishing accessibility in the distal regions during the reproductive process remain largely unclear. Our findings show that the SAS complex is recruited to numerous genes involved in flower development during the reproductive process and plays a crucial role in establishing their accessibility. While BAS and SAS target similar genomic regions, SAS exhibits a greater ability to regulate the accessibility of distal accessible regions. This suggests that the impact of SAS on chromatin accessibility is not only regulated by its binding to chromatin but also by its chromatin remodeling activity. Notably, distal accessible regions generally exhibit lower levels of histone acetylation compared to proximal promoter regions (Yan et al., 2019), indicating that differential histone modifications may regulate the remodeling activities of BAS and SAS. These findings are consistent with studies conducted in mammals, which have demonstrated the impact of histone modifications on the genomic targeting and remodeling activity of various SWI/SNF complexes (Mashtalir et al., 2021). In contrast to SAS and BAS, MAS primarily contributes to the accessibility of genes unrelated to flower development (such as housekeeping genes), further highlighting the functional specificities of different SWI/SNF complexes in Arabidopsis.

In mammals, there are three classes of SWI/SNF complexes: BAF, PBAF, and ncBAF. Among these, the BAF complex has been shown to specifically target distal enhancer regions, which play a crucial role in facilitating the transcription of genes involved in development (Alver et al., 2017). Similarly, in Arabidopsis, distal accessible regions have been identified as predictive markers for intergenic enhancers, exhibiting highly stage-specific patterns during flower development (Zhu et al., 2015; Yan et al., 2019). In this study, we present evidence that the SAS complex specifically contributes to the accessibility of distal accessible regions for numerous genes involved in flower development. This suggests that the SAS complex serves as the primary chromatin remodeler responsible for establishing the stage-specific accessibility of genes involved in flower development. Thus, it is likely that the SAS complex functions as the counterpart of the human BAF complex in regulating the accessibility of distal enhancer regions in genes related to flower development.

Transcription factors are known to have cell-type-specific functions, while chromatin remodeling complexes are generally considered to have broader regulatory roles. It is likely that the SAS complex collaborates with transcription factors to regulate gene transcription during reproductive development. Transcription factors can either bind the accessible cis-regulatory regions generated by chromatin remodelers or function as the pioneer transcription factors to bind the nucleosome DNA and recruitment the chromatin remodelers to generate accessibility. In Arabidopsis, LFY has been identified as a pioneer transcription factor that binds to nucleosome DNA and recruit SWI/SNF components to establish the accessibility of genes responsible for determining the floral organ identity (Jin et al., 2021; Lai et al., 2021). Certain MADS transcription factors, such as AP1 and SEP3, have also been suggested as potential pioneer factors involved in flower development (Lai et al., 2018). However, our results suggest that AP1 is not likely to be a pioneer transcription factor. Oppositely, the chromatin accessibility generated by SAS is crucial for the binding of AP1, indicating that AP1 functions downstream of chromatin accessibility. Interestingly, we found most of the accessible regions generated by SAS in inflorescences are occupied by AP1 with high enrichment, indicating that the chromatin remodeling of SAS is highly correlated with the flower development regulation of AP1 in inflorescences. This result is consistent with our motif analysis that MADS TF motifs are significantly enriched in the inflorescence-specific binding regions of SAS.

In mammals, the abundance of chromatin remodelers compared to transcription factors has facilitated the identification of chromatin remodelers co-purified with transcription factors. However, the identification of transcription factors co-purified with chromatin remodelers has proven challenging (Gourisankar et al., 2023). Similar difficulties have been encountered in plants, where MADS transcription factors have been used as baits to successfully identify SWI/SNF components, while attempts to identify MADS transcription factors using SWI/SNF components as baits have been unsuccessful (Smaczniak et al., 2012; Guo et al., 2022). In a recent online study, the interaction between transcription factors and components of chromatin-remodeling complexes during stomatal development was identified using proximity labeling methods (Liu et al., 2023). In the future, investigating the specific interaction between SWI/SNF chromatin complexes and different transcription factors during flower development will contribute to our understanding of the molecular mechanisms underlying the specific functions of the SAS complex in flower development.

## Materials and methods

### Plant materials

Arabidopsis were in the Columbia-0 (Col-0) ecotype. SWI/SNF mutants related to this study include: *brm-1* (SALK_030046C), *syd* (CS822017), *swi3d* (SALK_100310), *minu1* (SALK_015562), *minu2* (SALK_057856), *pms2a* (SALK_141512), *pms2b* (SALK_010411), and *sys1/2/3* (generated by CRISPR-Cas9 system), *ap1* (SALK_151561). Double mutants are generated by crossing. Detailed information on genotyping primers and CRISPR-Cas9 targets for SWI/SNF mutants can be found in the previous study (Guo et al., 2022). Full length genomic sequences of genes driven by 1.5-2 kb native promoter were cloned into modified pCAMBIA-1305-GFP or pCAMBIA-1305-3×Flag to generate C-terminal tagged transgenes for ChIP-seq. The primers used for these constructions can be found in the previous study (Guo et al., 2022). Arabidopsis seeds were sown on the Murahsige and Skoog (MS) medium and stratified at 4 ℃ for 2 days. Subsequently, they were grown under long-day conditions (16 h of light at 23 ℃ and 8 h of darkness at 22 ℃). Whole plants of 12-day-old seedlings grown on MS medium and inflorescences of 4∼5-week-old plants in soil were used for RNA-seq, ATAC-seq, and ChIP-seq experiments.

### RNA-seq and data analysis

About 0.1 gram of the inflorescence tips from the SWI/SNF mutants and the wild-type plants were harvested and ground in liquid nitrogen. For each genotype, three independent samples were collected as biological replicates. RNA was isolated by TRIzol reagent (Invitrogen, 15596018). Library construction and sequencing were carried out by BGI (Wuhan, China) using the DNBSEQ platform (sequencing method: SE50). The raw reads of RNA-seq were processed with the removal of adaptor sequences and the filter of low-quality reads. The clean reads were then mapped to the Arabidopsis genome using HISAT2 (version 2.2.1) (Kim et al., 2015) with default parameters. After counting the reads mapped on the exon by featureCounts (version 2.0.1)(Liao et al., 2014), edgeR (version 3.32.1) (Robinson et al., 2010) was used to evaluate the expression level, fold change and False Discovery Rate (FDR) of each gene. DEGs were filtered based on the criteria of |log2FC| ≥1 and FDR < 0.05.

### ATAC-seq and data analysis

About 0.1-0.2 gram of the inflorescence tips from the SWI/SNF mutants and the wild-type plants were used for ATAC-seq. Two biological replicates were set for each genotype. The ATAC-seq experiments were performed according to the previous study (Guo et al., 2022). After extracting and assessing the quality of the nuclei, about 10000∼50000 nuclei of each sample were collected for Tn5 tagmentation-based library construction using TruePrep DNA library Prep Kit V2 for Illumina (Vazyme, TD501). The ATAC-seq libraries were sent to Novogene and sequenced by an Illumina-NovaSeq 6000 (sequencing method: PE150).

The raw data were processed with Trim Galore (version 0.6.6) (Krueger, 2015) to remove adaptors and low-quality reads. Clean reads were then mapped to the Arabidopsis genome using bowtie2 (version 2.4.2) (Langmead and Salzberg, 2012) with the multi-mapped reads or PCR duplicates filtered out. The unique mapped reads were converted to bigwig files by bamCoverage in Deeptools (version 3.5.1)(Ramírez et al., 2016) with the parameter ‘binSize 10’ and ‘normalizeUsing RPKM’ and then visualized by the Integrative Genomics Viewer (IGV) (version 2.4.13) for snapshots. Accessible peaks were called using macs2 (version 2.2.7.1) (Zhang et al., 2008) with the following parameter: macs2 callpeak -t sample.bam -f BAMPE -g 1.2e+8 -nomodel --bdg --keep-dup all. Only the peaks that present in both replicates (irreproducible discovery rate < = 0.05) were retained for further analysis. Differentially accessible regions between the two samples were identified using the method ‘DESEQ2’ in DiffBind (version 3.0.15) (Rossinnes et al., 2011), with criteria of |log2FC| ≥0.5 and FDR < 0.05. Peaks were assigned to genes with the same rules as described in the previous study (Guo et al., 2022). Briefly, a peak that overlapped with the gene body including 1 kb upstream region were annotated to the corresponding gene, and if a peak was overlapped with more than one gene, it was annotated to the gene harboring the closest TSS to the midpoint of the peak.

### GSEA and GO analyses

The GSEA analysis was performed based on the gene sets generated from ATAC-seq analysis in this study and the RNA-seq data of various Arabidopsis tissues from the published study (Mergner et al., 2020). The expression level in flower and the average expression level across different tissues of the total Arabidopsis genes were used as the input expression dataset and the GSEA analysis was performed using GSEA software (version 4.3.2) according to the manual (Subramanian et al., 2005). The parameter ‘Metric for ranking genes’ was set as ‘Diff_of_Classes’. The term ‘flower specificity’ represents the expression level of a gene in the flower minus the average expression level of that gene across all the tissues. The GO analysis was performed by ClusterProfiler (version 4.2.2) (Yu et al., 2012) using the ‘org.At.tair.db’ database.

### ChIP-seq and data analysis

ChIP-seq were performed following the protocol from the previous studies with minor changes (Gendrel et al., 2005; Guo et al., 2022). Briefly, 4 grams of fresh seedlings or the inflorescence tips were collected and cross-linked in the buffer containing 1% formaldehyde under vacuum. The nuclei were extracted, washed, and sonicated by the Bioruptor 2000 in the sonication buffer containing 0.33% SDS. Subsequently, the fragmented chromatin was separated by centrifugation and the SDS concentration was diluted to 0.1%. For each sample, 0.1% of the total volume was set aside as the input sample, while the remaining diluted chromatin fragments were incubated overnight with the GFP antibody (Abcam, ab290) at a 1:600 dilution. The Dynabeads Protein A (Invitrogen, 100.01D) were added into the solution and incubated for 2-3 h, and then washed sequentially with low-salt buffer, high-salt buffer, and LiCl buffer, followed by two washes with TE buffer. The DNA-protein complexes were then eluted from the beads and de-crosslinked at 65℃ overnight. After the digestion of RNA and protein, the DNA was purified for library construction using NEBnext**®** Ultra^TM^ II DNA Library Prep Kit for Illumina (E7654S). The ChIP-seq libraries were sent to Novogene (Tianjin, China) for sequencing using an Illumina-NovaSeq 6000 (sequencing method: PE150).

After removal of adaptors from the raw reads, the clean reads were mapped to the Arabidopsis genome using Bowtie2 (version 2.4.2) with the same mapping method as for the ATAC-seq data. The binding peaks were called by macs2 (version 2.2.7.1) with the following parameters: macs2 callpeak -c input.bam -t IP.bam -f BAMPE -g 1.2e+8. Peaks were filtered and visualized using the same methods and standards as described in the ATAC-seq data analysis. Differential binding peaks of SWI/SNF subunits between seedlings and inflorescences were identified using the ‘DESEQ2’ method in DiffBind (version 3.0.15), with criteria of |log2FC| ≥0.5 and FDR < 0.05. The TF motif analysis was conducted by the findMotifsGenome.pl program in Homer (version 4.11)(Heinz et al., 2010) with default parameters using different sets of regions as input files.

### Plotting methods

The MDS plot were generated using the plotMDS function of the R package edgeR (version 3.32.1) with default parameters. The scatter plot was created using the ggscatter function of the R package ggpubr (version 0.4.0)(Kassambara, 2020). The PCA plot were generated using DiffBind (version 3.0.15) with default parameters. The heatmap based on Pearson correlation coefficients was generated using the heatmap.2 function of the R package gplots (version 3.1.1)(Warnes et al., 2014). Bar charts and the pie chart were created using Graphpad Prism (version 8.0). Box plots were generated using the boxplot function in R (version 4.1.0). The heatmap representing the enrichment of TF motifs was generated using Pheatmap (version 1.0.12) (Kolde, 2012). Venn diagrams, butterfly bar plots, the volcano plot, and the bubble plot containing multiple GO results of different samples were created using an online platform for data analysis and visualization at https://www.bioinfomatics.com.cn. The bubble plot showing the overlap of TF binding regions and the SAS-specific binding regions were plotted using ggplot2 (version 3.3.6).

### Accession numbers

The accession numbers of genes reported in this study are as follows: AT2G46020 (*BRM*), AT2G28290 (*SYD*), AT4G34430 (*SWI3D*), AT5G07940 (*SYS1*), AT5G07970 (*SYS2*), AT5G07980 (*SYS3*), AT3G06010 (*MINU1*), AT5G19310 (*MINU2*), AT3G08020 (*PMS2A*), AT3G52100 (*PMS2B*) and AT1G69120 (*AP1*).

## Data availability

The raw data for ATAC-seq, ChIP-seq, and RNA-seq experiments have been deposited in the Gene Expression Omnibus (GEO) database under the accession numbers GSE254654, GSE254655, and GSE254656, respectively. The access tokens are as follows: inapmusabpcvvar for GSE254654, wbahscwojrotpqn for GSE254655, and kfqbwguylhudhmh for GSE254656.

## Author contribution

JG and XJH conceived the project and designed the experiments. JG and ZZL performed the experiments. JG and YNS performed the bioinformatic analysis. JG and XJH wrote the manuscript.

## Acknowledgements

This work was supported by the National Natural Science Foundation of China to X. J. He (32025003).

## Supplemental information

**Supplemental Figure S1.**
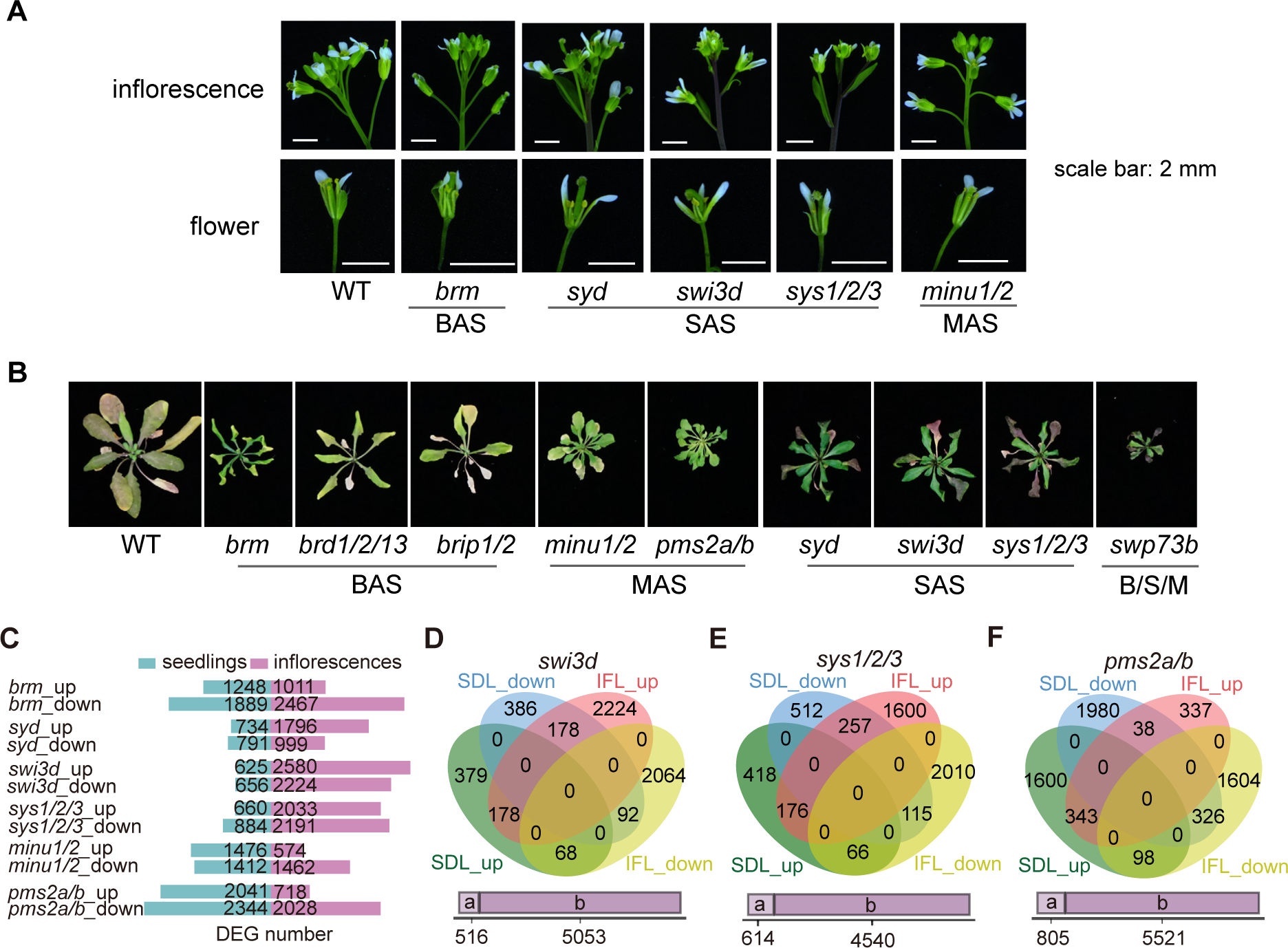
Inflorescence phenotype and different gene expression analysis in SWI/SNF mutants. **A)** Photographs of inflorescences and flowers from 5∼6-week-old SWI/SNF mutants. To show the inner stamens and pistil of a flower, some sepals and petals are removed in the second row of pictures. **B)** Images of rosette leaves from 8-week-old SWI/SNF mutants. To show the anthocyanin accumulation in rosette leaves, shoots and inflorescences were removed from the plant. **C)** The number of up-regulated and down-regulated DEGs in seedlings of inflorescences of SWI/SNF mutants, as determined by RNA-seq. **D-F)** Venn diagrams showing the overlap of up-regulated DEGs and down-regulated DEGs in seedlings and inflorescences of *swi3d* (**D**), *sys1/2/3* (**E**), and *pms2a/b* (**F**). The purple bar below the venn diagram marked by ‘a’ and ‘b’ represent genes shared by other sets and genes specifically belonging to one set of the venn diagrams, respectively.

**Supplemental Figure S2.**
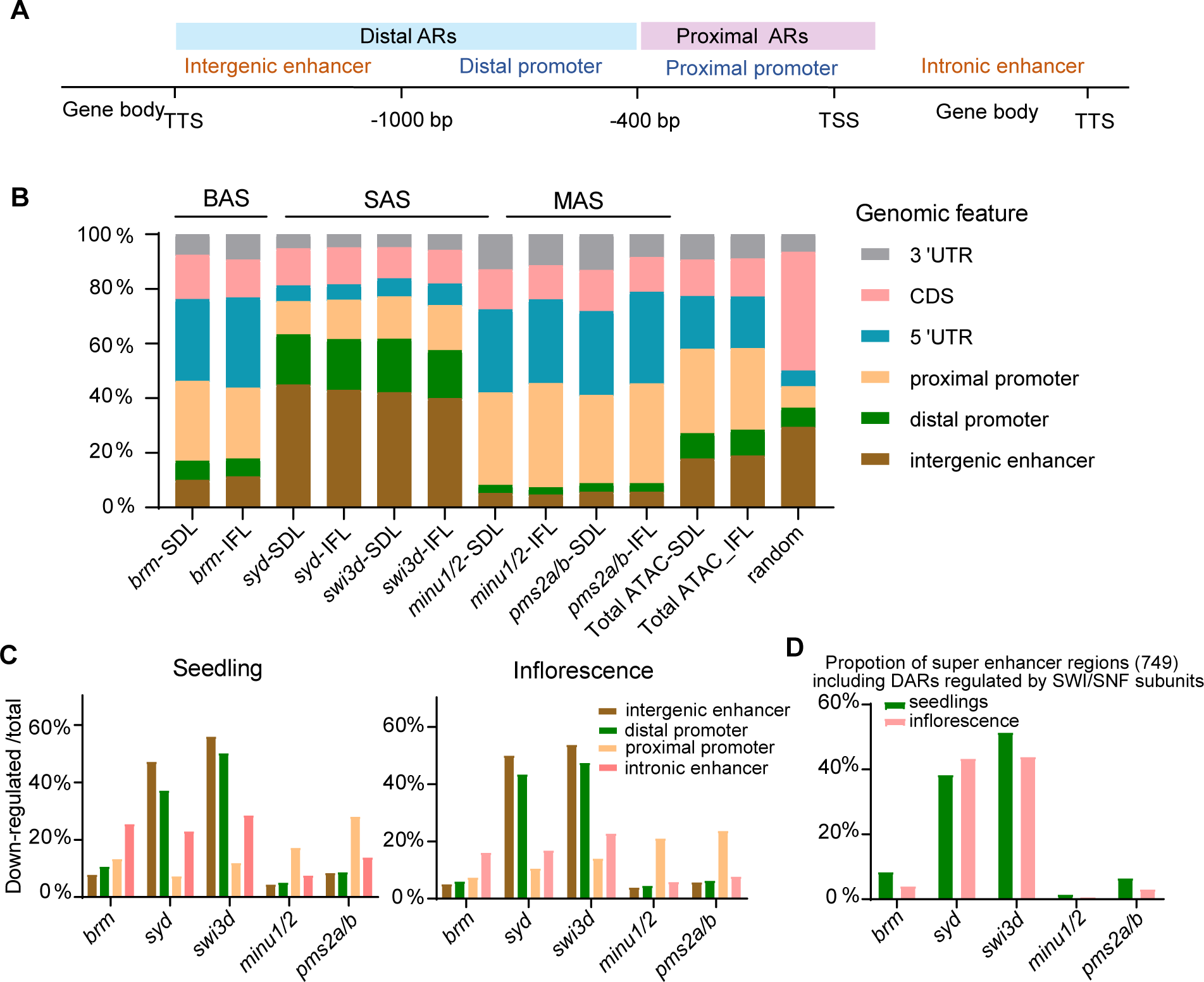
Proximal and distal accessible regions regulated by SWI/SNF complexes in seedlings and in inflorescences. **A)** The illustration of specified genomic features used for the peak annotation. ARs, accessible regions. **B)** Proportion of down-regulated differentially accessible regions annotated to specified features to the total down-regulated differentially accessible regions in seedlings or inflorescences of the indicated SWI/SNF mutants. The proportion of accessible regions annotated to specified genomic features to total accessible regions identified by ATAC-seq in the seedlings and inflorescences of wild-type plants is also presented. Random: 8,000 random genomic sites. DARs, differentially accessible regions. **C)** Proportion of downregulated differentially accessible regions in each SWI/SNF mutant to the total accessible regions annotated to specified genomic features in seedlings and in inflorescences. **D)** The proportion of super enhancers including accessible regions regulated by SWI/SNF components in seedlings and in inflorescences to total super enhancers identified in a previous study (Zhao et al., 2022).

**Supplemental Figure S3.**
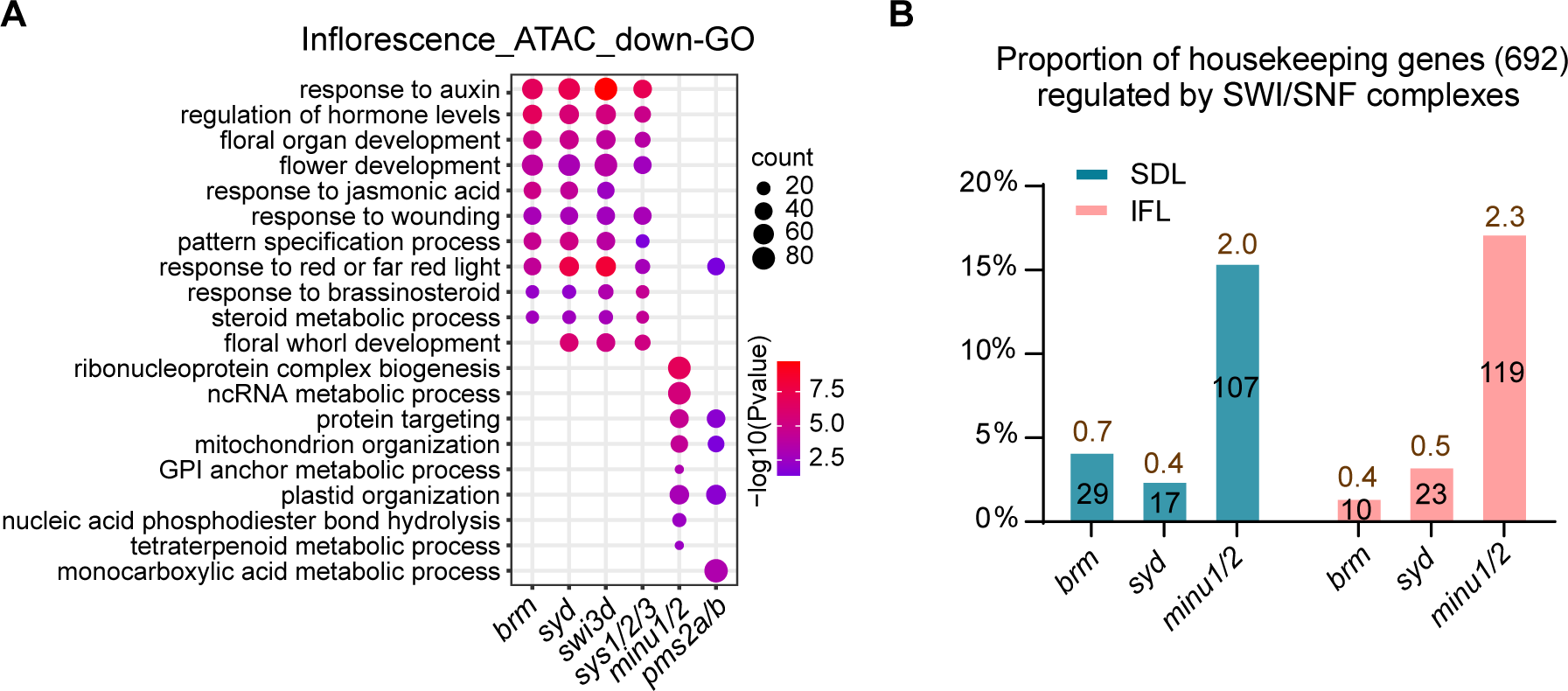
Characteristics of genes with down-regulated accessibility in inflorescences of SWI/SNF mutants. **A)** GO enrichment of genes with down-regulated accessibility in inflorescences of SWI/SNF mutants. Bubble plots show the enriched GO terms in genes with down-regulated accessibility in inflorescences of SWI/SNF muants. **B)** The propotions of housekeeping genes (identified by Cheng et al., 2017) showing down-regulated accessibility in seedlings (SDL) and inflorescences (IFL) of the SWI/SNF mutants. The number of overlaping genes are marked on the bars with black font. The values marked by the brown font above the bars represent the Representative Factors (the number of overlaping genes divided by the expected number of overlaping genes drawn from two independent groups).

**Supplemental Figure S4.**
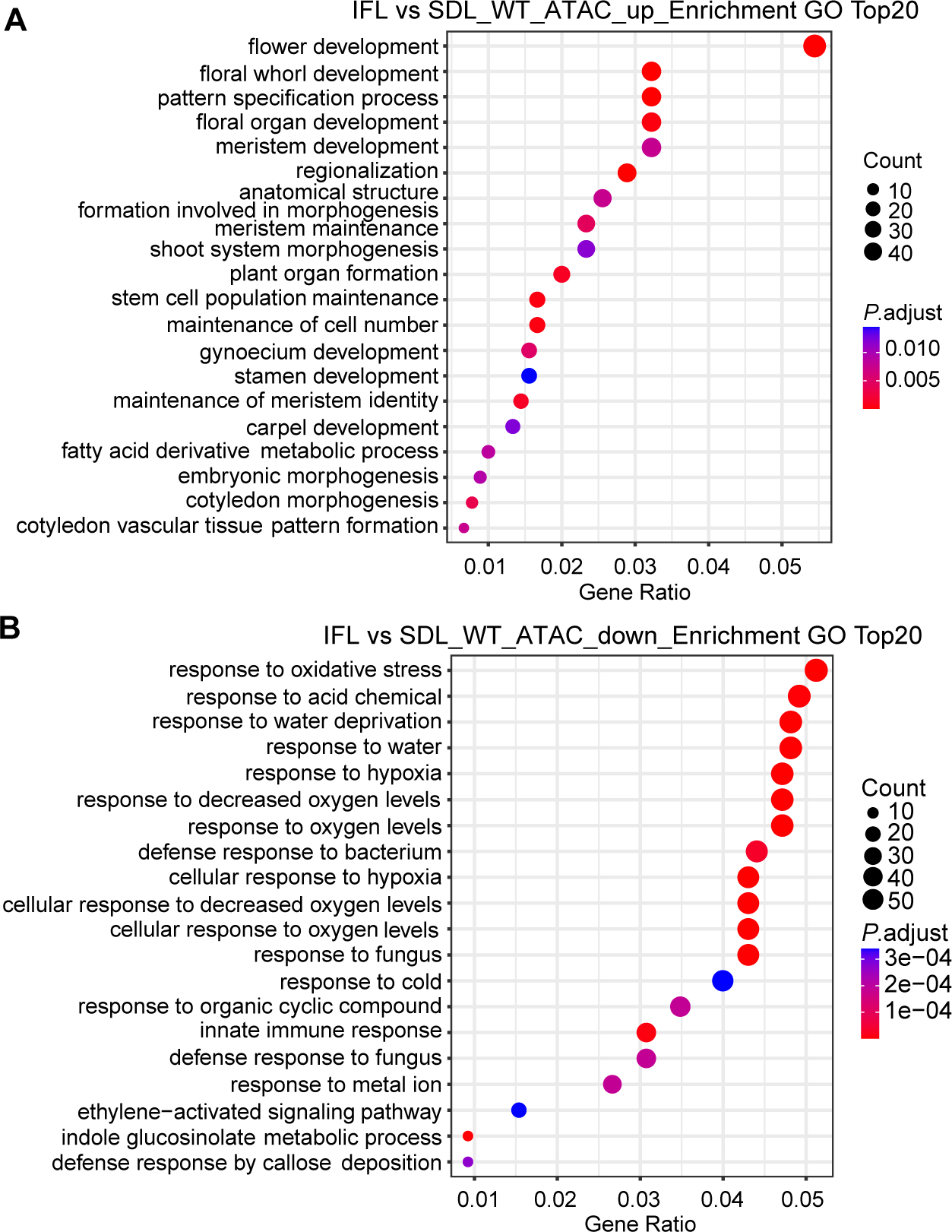
GO enrichment of seedling-specific and inflorescence-specific accessible genes. **A, B)** Bubble plots showing the top 20 enriched GO terms in specific accessible genes in inflorescences (**A**) and seedlings (**B**) of wild-type plants.

**Supplemental Figure S5.**
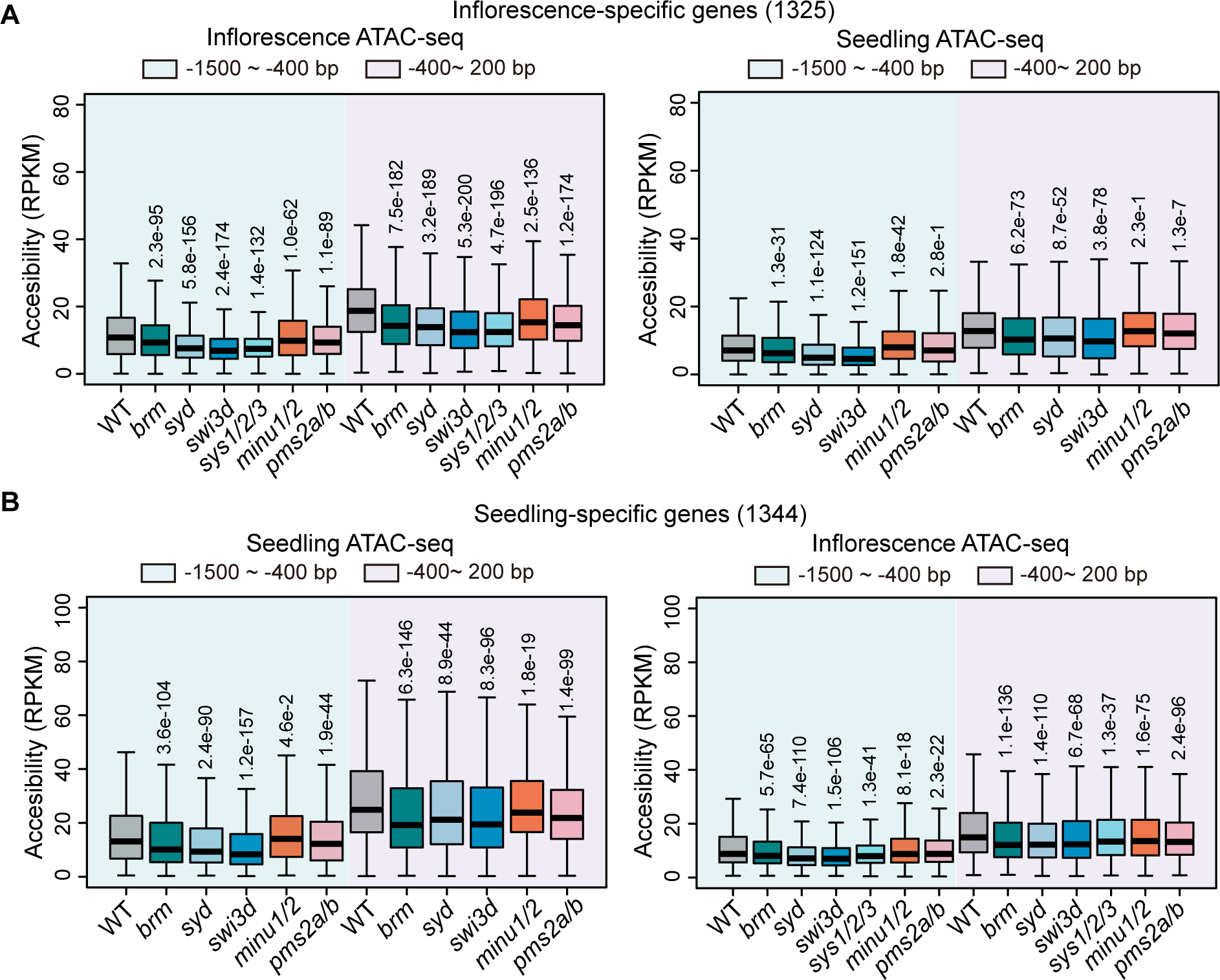
Effects of SWI/SNF subunit mutations on chromatin accessibility at inflorescence-specific accessible genes and seedling-specific genes. **A, B)** Boxplots showing the chromatin accessibility levels in seedlings and inflorescences of the wild type and SWI/SNF mutants at the 400-1500 upstream region of TSS and at the -400-200 bp TSS-flanking region. Analysis was conducted separately for inflorescence-specific genes (**A**) and seedling-specific genes (**B**). The center lines and box edges of the box plots represents medians and the interquartile range (IQR), respectively. Whiskers extend within 1.5 times the IQR. *P* values were determined by two-tailed Wilcoxon signed rank test between the mutants and the wild-type control in each group.

**Supplemental Figure S6.**
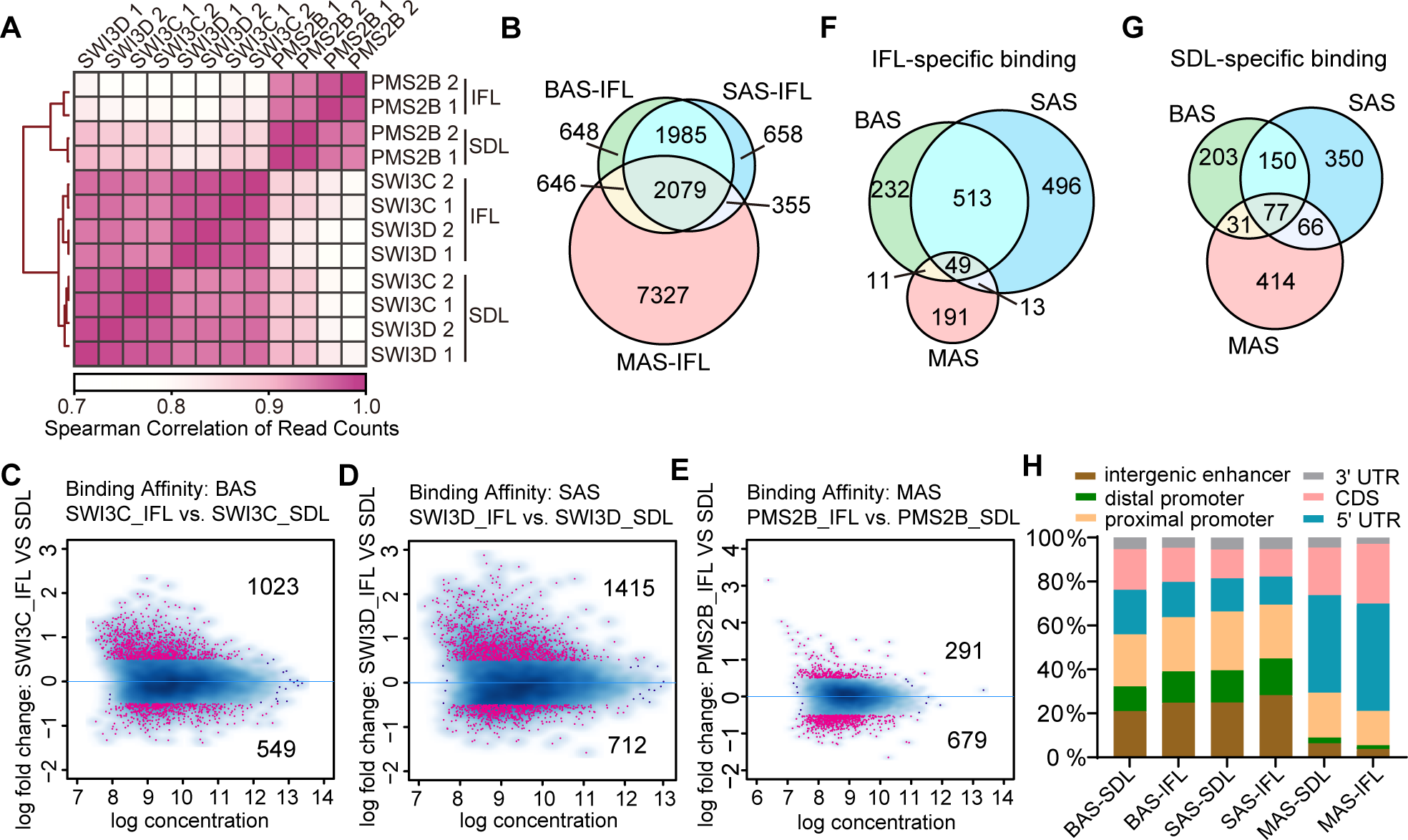
Comparison of the binding regions of BAS, SAS and MAS in seedlings and inflorescences. **A)** Heat map showing the pairwise Spearman correlation coefficients based on the ChIP-seq read counts of the two replicates of BAS (SWI3C), SAS (SWI3D), and MAS (PMS2B) subunits in seedlings or in inflorescences. **B)** Venn diagram illustrating the overlap of the genes bound by BAS, SAS and MAS in inflorescences. **C-E)** MA plots showing the up-regulated and down-regulated binding peaks of the BAS (**C**), SAS (**D**) and MAS (**E**) subunits in inflorescences compared to seedlings. The red points represent significant changed peaks with FDR<0.05 and |log2 (fold change) | >= 0.5. **F, G)** Venn diagrams showing the overlap of genes specifically bound by BAS, SAS and MAS in inflorescences (**F**) and in seedlings (**G**). **H)** The proportion of regions bound by BAS, SAS and MAS annotated to specified chromatin features in seedlings and in inflorescences. Random: 8,000 random genomic sites.

**Supplemental Figure S7.**
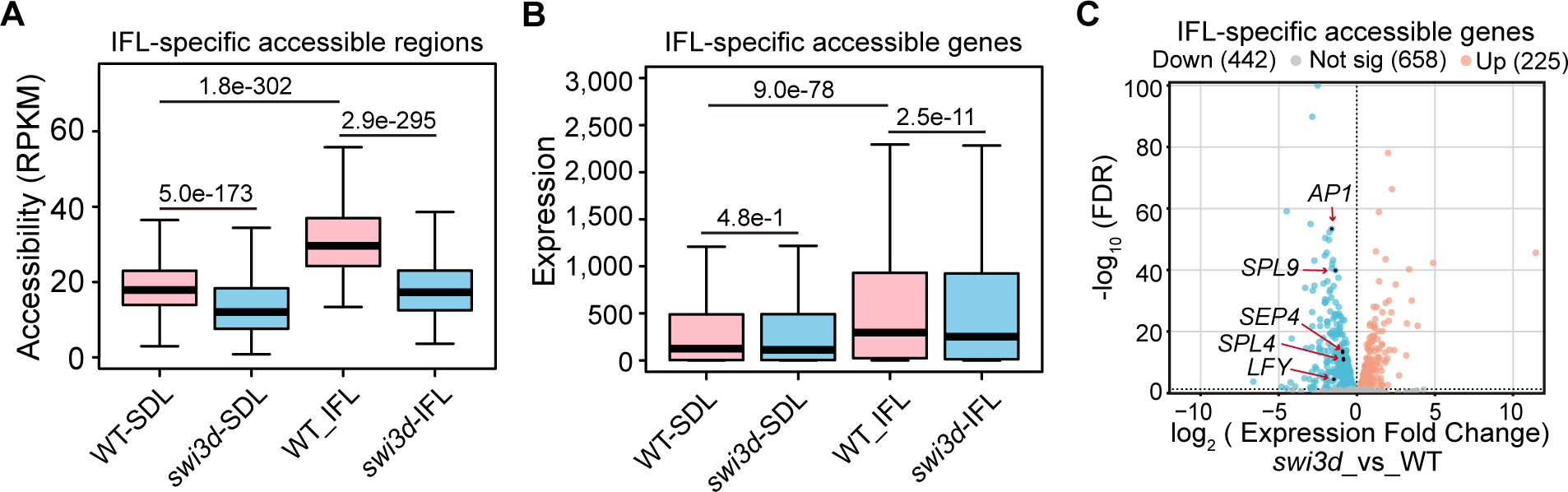
Effects of *swi3d* mutation on accessibility and expression of inflorescence-specific accessible genes. **A)** Boxplots showing the accessibility of inflorescence-specific accessible regions in seedlings and inflorescences of the wild type and *swi3d* mutant. **B)** Boxplots displaying the expression levels of inflorescence-specific accessible genes in seedlings and inflorescences of the wild type and *swi3d* mutant. The center lines and box edges of the box plots represent medians and the interquartile range (IQR), respectively. Whiskers extend within 1.5 times the IQR. *P* values were determined by two-tailed Wilcoxon signed rank test. **C)** Volcano plot showing the differential expression of inflorescence-specific accessible genes in the *swi3d* mutant compared to the wild type. The fold change and FDR were calculated using the inflorescence RNA-seq data. Genes that are up-regulated in *swi3d* (log2(fold change)>0, FDR<0.05) are marked in red, genes that are down-regulated (log2(fold change)<0, FDR<0.05) are marked in blue, and genes that are not significantly differentially expressed (FDR<0.05) are marked in grey. Representative points are marked with gene names.

**Supplemental Dataset 1.** Differentially expressed genes identified by RNA-seq in SWI/SNF mutants relative to wild type in inflorescences.

**Supplemental Dataset 2.** Differentially accessible regions of SWI/SNF mutants identified by ATAC-seq in inflorescences.

**Supplemental Dataset 3.** GO analysis of genes with decreased accessibility in SWI/SNF mutants identified by ATAC-seq.

**Supplemental Dataset 4.** Differentially accessible regions of wild-type plants between inflorescences and seedlings.

**Supplemental Dataset 5.** ChIP-seq peaks of SWI/SNF subunits in inflorescences and seedlings.

**Supplemental Dataset 6.** Differentially binding regions of SWI/SNF subunits identified by ChIP-seq between inflorescence and seedlings.

**Supplemental Dataset 7.** Transcription factor-binding motif analysis.

